# Ketamine reverses chronic stress-induced behavioral changes via Ca^2+^-permeable AMPA receptors in mice

**DOI:** 10.1101/2024.10.07.616991

**Authors:** Joshua C. Flowers, Paige E. Vetter, McKennon J. Wiles, Seung Hyun Roh, Ellison R. Black, Evelina Bouckova, Madison H. Wustrau, Rahmi Lee, Sang-Hun Lee, Seonil Kim

**Author notes:** These authors contributed equally (Joshua C. Flowers, Paige E. Vetter, McKennon J. Wiles, Seung Hyun Roh, Ellison R. Black, Evelina Bouckova, Madison H. Wustrau). Conflict of interest statement: The authors declare no competing financial interests.

## Abstract

**Background and Purpose:** Chronic stress affects brain functions leading to the development of mental disorders like anxiety and depression, as well as cognitive decline and social dysfunction. Among many biological changes in chronically stressed brains, disruptions in AMPA Receptor (AMPAR)-mediated synaptic transmission in the hippocampus are associated with stress responses. We have revealed that low-dose ketamine rapidly induces the expression of GluA1-containing, GluA2-lacking Ca^2+^-Permeable AMPARs (CP-AMPARs), which enhances glutamatergic synaptic strength in hippocampal neurons. Additionally, subanesthetic low-dose ketamine decreases anxiety-and depression-like behaviors in naïve animals. In addition to reducing depression, some research indicates that ketamine may have protective effects against chronic stress in both humans and animals. However, the role of CP-AMPARs in the actions of ketamine’s antistress effects is largely unknown.

**Experimental Approach:** We use whole-cell patch-clamp recordings from CA1 pyramidal neurons in female and male hippocampal slices and multiple behavioral assays including reciprocal social interaction, contextual fear conditioning, and tail suspension test.

**Key Results:** We demonstrate that low-dose ketamine treatment reverses chronic restraint stress (CRS)-induced social dysfunction, hippocampus-dependent fear memory loss, and depression-like behavior in both female and male mice. Furthermore, we show that the ketamine-induced antistress effects on these behaviors are dependent on CP-AMPARs.

**Conclusion and Implications:** Our findings suggest that subanesthetic low-dose ketamine rapidly triggers synaptic insertion of CP-AMPARs in the hippocampus, which induces antidepressant and antistress effects.

**What is already known:**

- After demonstrating rapid and robust antidepressant efficacy, the US Food and Drug Administration (FDA) approved esketamine (the S enantiomer form of ketamine) for the treatment of depression in 2019, sparking a surge in clinical and public interest around the world.
- Although the exact mechanisms are largely unknown, one mechanism postulated for ketamine’s actions involves the enhancement of excitatory synaptic drive through an increase in AMPA receptor (AMPAR)-mediated synaptic activity.

**What this study adds:**

- This study discovers that a single dose of ketamine rapidly (∼one hour) induces synaptic insertion of GluA1-containing, GluA2-lacking <u>C</u>a^2+^-<u>P</u>ermeable <u>AMPAR</u>s (CP-AMPARs), which is required for the antistress effects in animals.

**Clinical significance:**

- Ketamine is typically given in just a single dose or short series of doses rather than repeated treatments because of potential misuse and addictive properties. Furthermore, rapid relapse of depression occurs after cessation of ketamine treatment (18 - 19 days post-infusion).
- Understanding the precise mechanisms behind ketamine’s antidepressant effects could lead to the development of new clinical approaches to overcome its primary therapeutic limitation of restricted patient use.
- Unfortunately, the potential sex differences in response to ketamine have been particularly understudied at this time although the hypersensitivity of females to ketamine’s effects at lower doses has been shown in several animals’ studies. Therefore, our findings may help develop tailored treatment and dosing in clinics.

## Introduction

The World Health Organization (WHO) estimates that over 300 million people worldwide have depression (4.4% of the global population), which makes depression the leading cause of disability worldwide. The US Food and Drug Administration (FDA) approved esketamine (the S enantiomer form of ketamine) for the treatment of depression in 2019, sparking a surge in clinical and public interest around the world (Kohtala, 2021). Although the antidepressant effects of ketamine become evident within a few hours or 1 day of a single infusion, the benefits generally disappear within 1 week (Newport et al., 2015). More recent studies have shown that repeated ketamine infusions somewhat sustain the effects (aan het Rot et al., 2010; Murrough et al., 2013; J. L. Phillips et al., 2019). However, relapse of depression still occurs after cessation of treatment (on average, 18–19 days post-treatment) (aan het Rot et al., 2010; Murrough et al., 2013). Moreover, because ketamine is associated with psychomimetic side effects and risk of addiction, it is typically given in just a single dose or short series of doses rather than repeated treatments (Ng, Tse, Ng, & Lau, 2010; Sepulveda Ramos et al., 2022; Silva & Proulx, 2024). Some patients receiving ketamine therapy are also advised to stay on other traditional antidepressants whilst receiving ketamine (Veraart et al., 2021). However, reports have shown that some patients have been inflicted with adverse effects mimicking psychosis following ketamine administration on top of their regular polydrug regimen (Correia-Melo, Silva, Araujo-de-Freitas, & Quarantini, 2017; Paul, Schaaff, Padberg, Moller, & Frodl, 2009; Szymkowicz, Finnegan, & Dale, 2013). It is unclear which medications (or set of medications) negatively interact with ketamine (Veraart et al., 2021). Additionally, ketamine is found to be effective in reducing suicidality in treatment-resistant depression patients (10-20% of the depression patients) (Serafini, Howland, Rovedi, Girardi, & Amore, 2014). For these patients, ketamine + other traditional antidepressants is likely ineffective. Given all these problems, the field requires the development of new strategies that will permit a reduction in the frequency of ketamine administration after response to acute treatment, with maintenance of its beneficial effects for its broader application (Papakostas, 2020; J. L. Phillips et al., 2019). Unfortunately, achieving these goals is considerably more challenging because the exact mechanisms behind ketamine’s antidepressant effects are still unclear. Importantly, ketamine-induced neuroplasticity has been proposed as a mechanism underlying its antidepressant effects (Wu, Savalia, & Kwan, 2021). However, still unclear, is when and how neuroplasticity is induced. Specifically, molecular and cellular processes of how ketamine induces neuroplasticity remain to be elucidated.

One mechanism postulated for ketamine’s rapid antidepressant actions involves the enhancement of excitatory synaptic drive in the hippocampus (Aleksandrova, Wang, & Phillips, 2020; Kavalali & Monteggia, 2020; Miller, Moran, & Hall, 2016). However, it has been demonstrated that ketamine is a noncompetitive NMDA receptor (NMDAR) antagonist that inhibits excitatory synaptic transmission (Anis, Berry, Burton, & Lodge, 1983). This poses a contradictory question of how ketamine enhances excitatory activity and synaptic Ca^2+^ signaling while blocking NMDARs. This can be explained by an increase in AMPA receptor (AMPAR)-mediated synaptic activity but the mechanisms are unknown (Aleksandrova, Phillips, & Wang, 2017). In addition to reducing depression, some research indicates that ketamine may have protective effects against chronic stress in both humans and animals (Brachman et al., 2016; Feder et al., 2014; Gill et al., 2021; E. H. Lee, Park, Kwon, & Han, 2021; J. H. Ma et al., 2019; L. Ma et al., 2022; Taylor et al., 2018; Y. Yang et al., 2018; Zhou et al., 2021). Chronic stress affects brain functions, resulting in the development of mental disorders like anxiety and depression, as well as cognitive decline and social dysfunction (daSilva et al., 2021; Kessler, 1997; Lupien, McEwen, Gunnar, & Heim, 2009). Among many biological changes in chronically stressed brains, disruptions in AMPAR-mediated synaptic transmission in the hippocampus are associated with stress responses in animal models and individuals with depression (He, Zhou, Wang, & Chen, 2023).

Thus, recovery from AMPAR-mediated synaptic dysfunction may be the key target of ketamine’s antistress and antidepressant effects.

There are two general types of AMPARs formed through combination of their subunits: GluA2-containing Ca^2+^-impermeable AMPARs and GluA2-lacking, GluA1-containing Ca^2+^-Permeable AMPARs (CP-AMPARs) (Diering & Huganir, 2018). Compared to Ca^2+^-impermeable AMPARs, CP-AMPARs have higher single channel conductance, which contributes to synaptic plasticity (Aoto, Nam, Poon, Ting, & Chen, 2008; Goel & Lee, 2007; Goel et al., 2011; S. Kim, Violette, & Ziff, 2015; S. Kim & Ziff, 2014; H. K. Lee, 2012; Purkey & Dell’Acqua, 2020; Sanderson, Gorski, & Dell’Acqua, 2016; Sanderson, Scott, & Dell’Acqua, 2018; Thiagarajan, Lindskog, & Tsien, 2005). Both preclinical and clinical studies show that chronic stress significantly reduces GluA1 levels in hippocampal synapses, while there are conflicting results describing alterations in hippocampal GluA2 under chronic stress (He et al., 2023). These results suggest that the stress-induced decrease in synaptic GluA1 levels diminishes excitatory synaptic transmission in the hippocampus and exacerbates negative behaviors in chronic stress. Therefore, recovery from GluA1-mediated synaptic dysfunction may be the key target of ketamine’s antidepressant and antistress effects. We have revealed that low-dose ketamine (1 μM, a concentration that blocks approximately 50% of NMDAR-induced currents) (Hare et al., 2019) rapidly induces the expression of CP-AMPARs in cultured hippocampal neurons, which is mediated by increasing GluA1 phosphorylation and its surface expression (Zaytseva et al., 2023). Moreover, we have shown that this ketamine-induced CP-AMPAR expression enhances glutamatergic synaptic strength in cultured neurons (Zaytseva et al., 2023). Additionally, subanesthetic low-dose ketamine (5-10 mg/kg) decreases anxiety-and depression-like behaviors in naïve animals (Zaytseva et al., 2023). In fact, an increase in hippocampal activity is known to improve anxiety-and depression-like behaviors, fear memory, and social behavior in mice (Aleksandrova et al., 2020; Bird & Burgess, 2008; Grieco et al., 2022; Kavalali & Monteggia, 2020; Miller et al., 2016; Okuyama, Kitamura, Roy, Itohara, & Tonegawa, 2016; Sun et al., 2020; Tovote, Fadok, & Luthi, 2015). This suggests that ketamine-induced recovery from disruptions in AMPAR-mediated synaptic transmission and behaviors in stressed animals is likely associated with an increase in synaptic GluA1 expression in the hippocampus (Fischell, Van Dyke, Kvarta, LeGates, & Thompson, 2015; Kallarackal et al., 2013; Li et al., 2011).

Although we have shown that subanesthetic low-dose ketamine selectively increases GluA1 expression in the hippocampus and induces antidepressant effects via CP-AMPAR expression in naïve mice (Zaytseva et al., 2023), much research with hippocampal neural circuits is required to fully comprehend the ketamine’s actions on chronic stress-induced behavioral alterations. Here, we use an electrophysiological approach in hippocampal slices to confirm that low-dose ketamine treatment indeed induces synaptic insertion of CP-AMPARs in hippocampal pyramidal neurons. We further show that ketamine at the low dosages reverses chronic restraint stress (CRS)-induced behavioral changes in mice, which is dependent on CP-AMPARs. Our findings thus suggest that subanesthetic low-dose ketamine rapidly triggers synaptic insertion of CP-AMPARs in the hippocampus, which in turn enhances synaptic strength and compensate for reduced NMDAR antagonism-mediated synaptic Ca^2+^ signaling to induce antidepressant and antistress effects.

## Methods

### Animals

C57Bl6J (Jax 000664) mice were obtained from Jackson laboratory and bred in the animal facility at Colorado State University (CSU). Animals were housed under a 12:12 hour light/dark cycle. To collect brain tissues from animals, mice were deeply anesthetized and euthanized by CO_2_ asphyxiation. All efforts were made to minimize animal suffering. CSU’s Institutional Animal Care and Use Committee (IACUC) reviewed and approved the animal care and protocol (3408). Results were reported following the ARRIVE (Animal Research: Reporting of In Vivo Experiments) guidelines (Kilkenny et al., 2010).

### Chronic Restraint stress (CRS)

3-month-old C57Bl6 female and male mice were individually placed into 50 mL polypropylene conical tubes with multiple holes for ventilation, and they were exposed to restraint stress (2LJhr/day) for 14 consecutive days. After restraint stress, mice were returned to the home cage. Following 2-week CRS, drug injections, blood collection, and multiple behavior assays were carried out as shown in the experimental timeline in **Fig. 1**.

**Figure 1.**
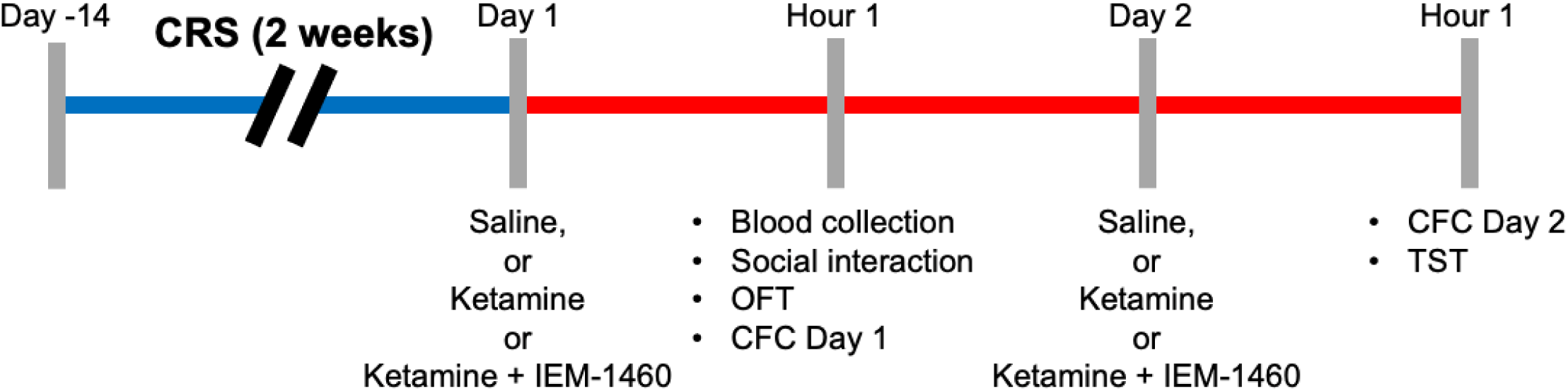
Experimental timeline of chronic restraint stress (CRS) that is applied to mice for 2 weeks, drug treatment, blood collection, and behavioral tests - reciprocal social interaction test, open-field test (OFT), contextual fear conditioning (CFC) day 1 and 2, and tail suspension test (TST). Single intraperitoneal injection of ketamine (5 mg/kg for females and 10 mg/kg for males) or saline in CRS mice in the presence or absence of IEM-1460 (10 mg/kg for both females and males). We optimized the test order from least to most stressful in order to prevent the possibility that multiple behavioral tests with the same mouse can induce stress, especially if the tests are performed too closely together.

### Drug treatment

Ketamine hydrochloride (VetOne, 510189) was used in both *in vitro* and *in vivo* experiments. 5 mg/kg and 10 mg/kg ketamine were intraperitoneally injected into 3-month-old female and male C57Bl6 mice, respectively (the conditions have been shown to exclusively increase hippocampal GluA1 and induce antidepressant effects on naïve animals’ behaviors) (Zaytseva et al., 2023), and saline was administered to controls. We have approval from IACUC to use ketamine and have the United States Drug Enforcement Administration license to use ketamine for research purpose (DEA# RK0573863). 100 μM 1-Naphthyl acetyl spermine trihydrochloride (NASPM, Tocris Bioscience) was used to block CP-AMPARs in slice electrophysiology. 10 mg/kg IEM-1460 (Tocris Bioscience, 1636) was intraperitoneally injected into 3-month-old male and female C57Bl6J mice to inhibit *in vivo* CP-AMPAR activity because it is blood-brain barrier (BBB)-permeable (Adotevi et al., 2020; Szczurowska & Mares, 2015; Wiltgen et al., 2010). Following CRS, we performed the experiments in the presence or absence of IEM-1460 and compared outcomes to examine whether ketamine treatment reversed the CRS-induced effects via CP-AMPARs.

### Brain slice preparation and electrophysiology

The mice were deeply anesthetized with isoflurane, and their brains were removed. The brains were submerged in cold, oxygenated (95% O_2_ and 5% CO_2_) slicing medium containing (in mM): 85 NaCl, 1.25 NaH_2_PO_4_, 4 MgCl_2_, 0.5 CaCl_2_, 24 NaHCO_3_, 2.5 KCl, 75 sucrose, and 25 glucose. Horizontal hippocampal slices (350 μm thick) were cut using a vibratome (Leica VT 1000S; Leica Microsystems). The solution was continuously oxygenated with 95% O_2_ and 5% CO_2_. Slices were initially maintained at 33°C for 30 min, then at room temperature (22 ± 1°C) until they were used for electrophysiological recordings. Whole-cell patch-clamp recording procedures used in the present study were like those described previously (Govindaiah et al., 2018; Kang et al., 2020; Kang et al., 2018). Electrophysiological recordings were performed on CA1 pyramidal cells at 33°C in oxygenated (95% O_2_ and 5% CO_2_) artificial cerebrospinal fluid (ACSF) containing (in mM): 126 NaCl, 2.5 KCl, 26 NaHCO_3_, 2 CaCl_2_, 2 MgCl_2_, 1.25 NaH_2_PO_4_, and 10 glucose. Slices were superfused with ACSF at a rate of 2.0 ml/min. Individual neurons were visualized using an upright microscope (Olympus BX51WI) with infrared differential interference contrast (DIC) optics. CA1 pyramidal cells in the stratum pyramidale (the pyramidal cell layer), visualized using differential interference contrast microscopy, will be selected based on their morphology as described previously (Kang et al., 2020). Whole-cell patch-clamp recordings were obtained from CA1 pyramidal cells with borosilicate patch pipettes (2–4 MΩ) filled with an internal solution containing (in mM): 126 K-gluconate, 4 KCl, 10 HEPES, 4 ATP-Mg, 0.3 GTP-Na, and 10 phosphocreatine. The pH and osmolarity of the internal solution were adjusted to 7.2 and 290 mOsm, respectively. Recordings were obtained using a MultiClamp 700B amplifier (Molecular Devices). After forming the whole-cell configuration, the recording was allowed to rest for at least 5 min prior to data acquisition. Signals were sampled at 10 kHz, low-pass filtered at 3 kHz using Digidata 1440A analog-to-digital digitizer (Molecular Devices) and stored on computer for subsequent analysis using pClamp 10 software (Molecular Devices). Series resistances were continuously monitored with recordings being discarded if the series resistance increased >20%. We then measured glutamate uncaging-induced excitatory postsynaptic currents (uEPSCs) in hippocampal CA1 neurons one hour following a single dose of ketamine (5 mg/kg for females and 10 mg/kg for males). For glutamate uncaging, 100 μM MNI-caged L-glutamate was added to ACSF, and epi-illumination photolysis (390 nm, 0.12 mW/mm^2^, 50 ms) was applied near the soma of a neuron. 1 μM TTX was added to ACSF for preventing action potential-dependent network activity. The average of uEPSC amplitudes (V_hold_ =-70 and +60 mV) in CA1 pyramidal cells was compared between saline and ketamine treatment. To examine whether CP-AMPARs played a role in uEPSCs, we recorded uEPSCs with a bath application of 100 μM NASPM, a CP-AMPAR blocker. Additionally, we added saline to ACSF as a control for NASPM treatment. Each recording was performed in different slices. We then analyzed the amplitudes of uEPSCs in each condition using Clampfit 10.7 (Molecular Devices).

### Reciprocal social interaction

We performed the reciprocal social interaction test to examine social interaction between two freely moving mice (Bolivar, Walters, & Phoenix, 2007; Matsuo et al., 2009). A test mouse (3-month-old) and a stranger mouse of identical genotypes that were previously housed in different cages were placed into the chamber (40 W x 40 L x 40 H cm) and allowed to explore freely for 20 min. Social behavior was recorded using a camera mounted overhead. Social contacts were determined by the sniffing time that was defined as each instance in which a test mouse’s nose came within 2 cm toward a stranger mouse. The total number of reciprocal interactions and total duration of reciprocal interactions were measured manually. Animals that escaped the chamber or showed aggressive behaviors (e.g. attacking a stranger mouse) were excluded from the analysis.

### Open field test (OFT)

We measured locomotor activity and anxiety-like behavior using OFT as carried out previously (Kang et al., 2021; Shou, Tran, Snyder, Bleem, & Kim, 2018; Zaytseva et al., 2023). A test mouse was first placed in the center of the open field chamber (40 W x 40 L x 40 H cm) for 5 min. Animals were then allowed to explore the chamber for 20 min. The behavior was recorded by a video camera. A 20 x 20 cm center square was defined as the inside zone. Data were analyzed using the ANY-maze tracking program (Stoelting Co.) to acquire total traveled distance (locomotor activity) and time spent outside and inside (anxiety-like behavior). Animals that escaped the chamber or showed significantly decreased locomotor activity were excluded from the analysis.

### Contextual fear conditioning (CFC)

CFC (Habitest Modular System, Coulbourn Instrument) was carried out as described previously (S. Kim et al., 2016; R. Lee, Kim, & Kim, 2024; Shou et al., 2018). On Day 1, a test mouse was placed in a novel rectangular chamber with a grid floor. After a 3-min baseline period, the test animal was given one shock (a 2 sec, 0.5 mA shock) and stayed in the chamber for an additional 1 min after the shock before being returned to the home cage overnight. A contextual memory test was conducted the next day (Day 2) in the same conditioning chamber for 3 min. Fear memory was determined by measuring the percentage of the freezing response (immobility excluding respiration and heartbeat) using an automated tracking program (FreezeFrame). Mice that did not freeze were excluded from the analysis.

### Tail suspension test (TST)

It has been suggested that the forced swim test (FST) and TST are based on the same principle: measurement of the duration of immobility when rodents are exposed to an inescapable situation (Castagne, Moser, Roux, & Porsolt, 2011). Importantly, TST is known to be more sensitive to antidepressant agents than FST because the animals remain immobile longer in TST than FST (Cryan, Mombereau, & Vassout, 2005). We thus used TST to examine depression-like behavior as described previously (S. Kim et al., 2016; S. Kim, Shou, Abera, & Ziff, 2018; Zaytseva et al., 2023). A test mouse was suspended by its tails from a rod suspended 20 cm above the tabletop surface with adhesive tape placed 1 cm from the tip of the tail. Animals were immobile when they exhibited no body movement and hung passively for > 3 sec. The time during which mice remained immobile was quantified over a period of 6 min. The behavior was recorded by a video camera and, where data were analyzed using the ANY-maze tracking program to acquire immobility (depression-like behavior).

Mice that successfully climbed their tails to escape were excluded from the analysis.

### Corticosterone

Corticosterone levels in mice were measured as described previously (Luo et al., 2016; McCosh, Kreisman, Tian, Thomas, & Breen Church, 2024). 50 µl tail-tip whole blood samples were collected and centrifuged at 5,000 rpm for 10 min. 10 µl of serum was harvested and stored at −20°C until assay. 5 µl of serum samples in each condition was assayed using the DetectX® Corticosterone Multi-Format Enzyme-linked immunosorbent assay (ELISA) kit (Arbor Assays, Inc. K014-H). To prevent possible artifacts during the ELISA experiment, we measured the corticosterone levels twice using the same sample.

### Statistical analysis

All behavior tests were blindly scored by more than two investigators. The Franklin A. Graybill Statistical Laboratory at Colorado State University was consulted for statistical analysis in the current study, including sample size determination, randomization, experiment conception and design, data analysis, and interpretation. We used the GraphPad Prism 10 software to determine statistical significance (set at *p* < 0.05). Grouped results of single comparisons were tested for normality with the Shapiro-Wilk normality or Kolmogorov-Smirnov test and analyzed using an unpaired two-tailed Student’s t-test when data are normally distributed. Differences between multiple groups were assessed by N-way analysis of variance (ANOVA) with the Tukey test. The graphs were presented as mean ± Standard Deviation (SD). We discard data points that are located further than two SD above or below the average as an outlier.

## Results

### Subanesthetic low-dose ketamine induces synaptic insertion of CP-AMPARs in hippocampal CA1 pyramidal neurons

The hippocampus is one of the key brain regions controlling cognition, motivation, and social behavior (Felix-Ortiz, Burgos-Robles, Bhagat, Leppla, & Tye, 2016; Hong & Kaang, 2022; W. B. Kim & Cho, 2020; M. L. Phillips, Robinson, & Pozzo-Miller, 2019; Schumacher, Vlassov, & Ito, 2016; Sun et al., 2020; Tao et al., 2022; X. Wang & Zhan, 2022; Watarai, Tao, Wang, & Okuyama, 2021; Yoshida et al., 2021). An increase in hippocampal activity reverses stress-induced memory impairment, social dysfunction, and mood disorder-associated behaviors (Aleksandrova et al., 2020; Bird & Burgess, 2008; Grieco et al., 2022; Kavalali & Monteggia, 2020; Miller et al., 2016; Okuyama et al., 2016; Su, Lu, Fu, Geng, & Chen, 2023; Sun et al., 2020; Tovote et al., 2015). More importantly, the hippocampus is preferably targeted by ketamine at the low doses (P. A. Davoudian, Shao, & Kwan, 2023). This suggests that the hippocampus is the primary brain region to be responsible for ketamine’s antistress effects. In fact, we have discovered that subanesthetic low-dose ketamine significantly increases synaptic GluA1 but not GluA2, an indication of CP-AMPAR expression, in the hippocampus of naïve animals (Zaytseva et al., 2023). We thus decided to examine whether low-dose ketamine was able to induce synaptic insertion of CP-AMPARs in hippocampal neurons using whole-cell patch clamping with glutamate uncaging. We intraperitoneally injected 5 mg/kg and 10 mg/kg ketamine to female and male C57Bl6 mice, respectively, and saline was administered to controls. These ketamine concentrations were shown to exclusively increase hippocampal GluA1 and induce antidepressant effects on naïve animals’ behaviors (Zaytseva et al., 2023). One hour following injection, hippocampal slices were obtained, and we then measured the amplitudes of glutamate uncaging-induced excitatory postsynaptic currents (uEPSCs) (V_hold_ = - 70 and +60 mV) in CA1 pyramidal cells. We found that ketamine injection significantly elevated the amplitudes of uEPSCs (V_hold_ =-70 mV) in these neurons when compared to control cells (CTRL) (FEMALE: CTRL, 86.330 ± 23.107 pA and Ketamine, 113.208 ± 30.563 pA, *p =* 0.0367. MALE: CTRL, 83.915 ± 27.768 pA and Ketamine, 109.329 ± 28.293 pA, *p* = 0.0471) (**Fig. 2a and Table 1**). However, when ketamine had no effect on uEPSCs (V_hold_ = +60 mV) (FEMALE: CTRL, 292.223 ± 97.252 pA and Ketamine, 360.238 ± 173.766 pA, *p =* 0.5017. MALE: CTRL, 286.115 ± 129.521 pA and Ketamine, 372.100 ± 126.540 pA, *p* = 0.2718) (**Fig. 2b and Table 2**). To examine whether CP-AMPARs were involved in the ketamine’s effects on uEPSCs, we added 100 μM NASPM, a CP-AMPAR antagonist, to ACSF and measured the amplitudes of uEPSCs in CA1 pyramidal cells from saline-or ketamine-injected animals. We found that NASPM treatment significantly reduced the amplitudes of uEPSCs (V_hold_ =-70 and + 60 mV) following ketamine treatment (V_hold_ =-70 mV; FEMALE: Ketamine + NASPM, 58.671 ± 22.786 pA, *p* < 0.0001. MALE: Ketamine + NASPM, 55.159 ± 26.805 pA, *p* < 0.0001. V_hold_ = +60 mV; FEMALE: Ketamine + NASPM, 213.793 ± 111.514 pA, *p* = 0.0170. MALE: Ketamine + NASPM, 233.635 ± 130.302 pA, *p* = 0.0133) (**Fig. 2a-2b and Table 1-2**). However, NASPM had no effect on the amplitudes of uEPSCs (V_hold_ =-70 and + 60 mV) measured in slices from saline-injected mice (V_hold_ =-70 mV; FEMALE: NASPM, 61.702 ± 21.324 pA, *p* = 0.0575. MALE: NASPM, 77.500 ± 16.628 pA, *p* = 0.9248. V_hold_ = +60 mV; FEMALE: NASPM, 276.986 ± 97.353 pA, *p* = 0.9885. MALE: NASPM, 284.958 ± 123.642 pA, *p* > 0.9999) (**Fig. 2a-2b and Table 1-2**). These findings demonstrate that subanesthetic low-dose ketamine induces synaptic insertion of CP-AMPARs in hippocampal CA1 pyramidal neurons.

**Figure 2.**
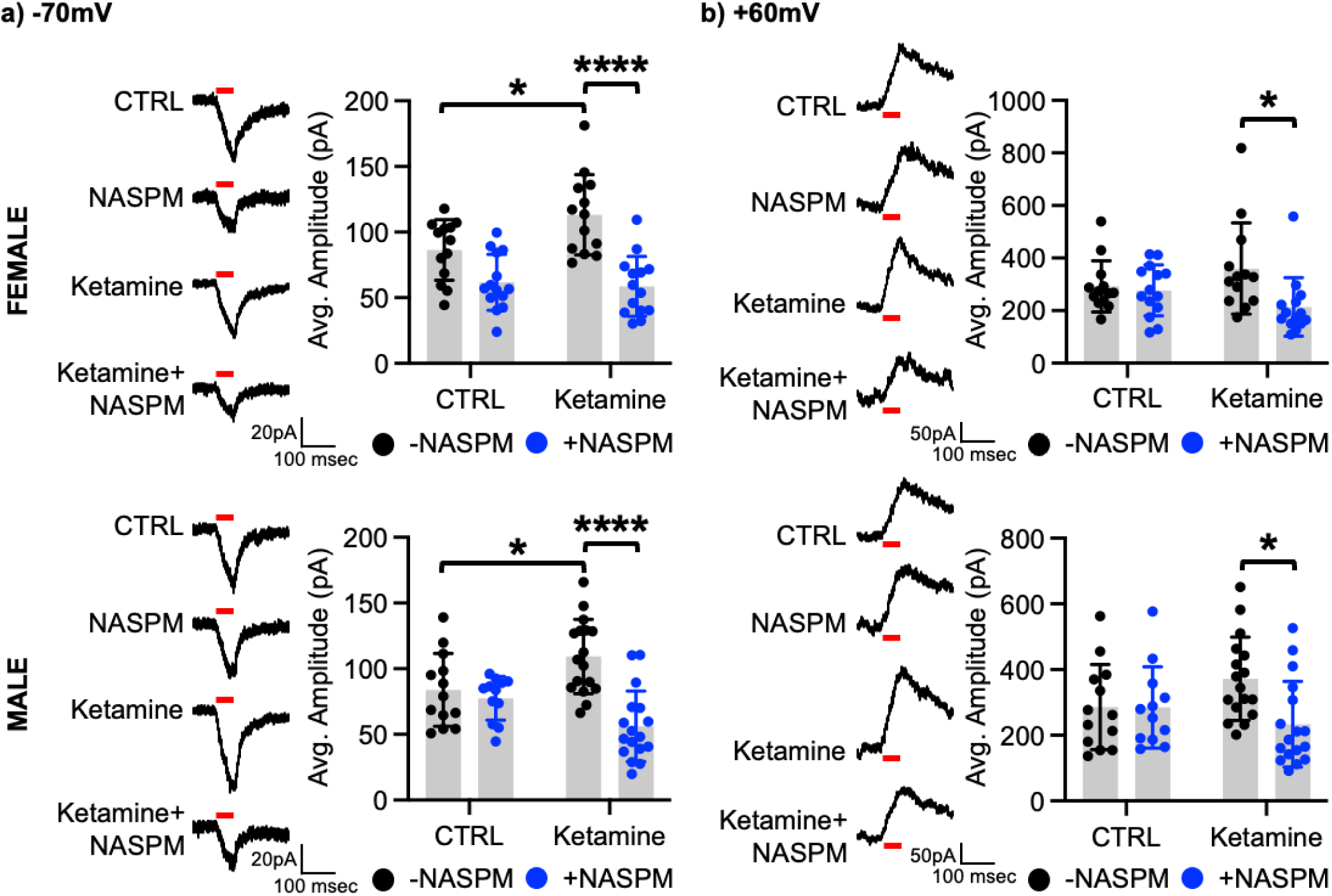
Subanesthetic low-dose ketamine induces synaptic insertion of CP-AMPARs in hippocampal CA1 pyramidal neurons. Representative traces and summary of the amplitudes of uEPSCs (**a)** V_hold_ =-70 mV and **b)** V_hold_ = +60 mV) in each condition (n = number of cells [n = number of animals]. FEMALE: CTRL = 13 [3], NASPM = 14 [3], Ketamine = 13 [3], and Ketamine + NASPM = 14 [3]. MALE: CTRL = 13 [4], NASPM = 14 [4], Ketamine = 13 [3], and Ketamine + NASPM = 14 [3]). \**p* < 0.05 and \*\*\*\**p* < 0.0001, Two-Way ANOVA, Tukey test. A bar indicates photostimulation. Ketamine: 5 mg/kg for females and 10 mg/kg for males. NASPM: 100 μM.

**Table 1.** Statistical analysis of the amplitudes of uEPSCs (V_hold_ =-70 mV) in hippocampal CA1 pyramidal neurons.

|  |  |  |  |  |  |
| --- | --- | --- | --- | --- | --- |
| Female (-70 mV) |  |  |  |  |  |
| ANOVA table | SS (Type III) | DF | MS | F (DFn, DFd) | P value |
| Interaction | 3015 | 1 | 3015 | F (1, 50) = 4.979 | P=0.0302 |
| Row Factor (Saline vs. Ket) | 1917 | 1 | 1917 | F (1, 50) = 3.165 | P=0.0813 |
| Column Factor (Saline vs. NASPM) | 21122 | 1 | 21122 | F (1, 50) = 34.88 | P<0.0001 |
| Residual | 30277 | 50 | 605.5 |  |  |
| Tukey's multiple comparisons test | Predicted (LS) mean diff. |  | 95.00% CI of diff. |  | Adjusted P Value |
| CTRL vs. NASPM | 24.63 |  | -0.5609 to 49.82 |  | 0.0575 |
| CTRL vs. Ketamine | -26.88 |  | -52.53 to -1.227 |  | 0.0367 |
| CTRL vs. Ketamine + NASPM | 27.66 |  | 2.470 to 52.85 |  | 0.0262 |
| NASPM vs. Ketamine | -51.51 |  | -76.69 to -26.32 |  | <0.0001 |
| NASPM vs. Ketamine + NASPM | 3.031 |  | -21.69 to 27.75 |  | 0.9879 |
| Ketamine vs. Ketamine + NASPM | 54.54 |  | 29.35 to 79.72 |  | <0.0001 |
| Male (-70 mV) |  |  |  |  |  |
| ANOVA table | SS (Type III) | DF | MS | F (DFn, DFd) | P value |
| Interaction | 7866 | 1 | 7866 | F (1, 55) = 11.82 | P=0.0011 |
| Row Factor (Saline vs. Ket) | 59.69 | 1 | 59.69 | F (1, 55) = 0.08970 | P=0.7657 |
| Column Factor (Saline vs. NASPM) | 12776 | 1 | 12776 | F (1, 55) = 19.20 | P<0.0001 |
| Residual | 36598 | 55 | 665.4 |  |  |
| Tukey's multiple comparisons test | Predicted (LS) mean diff. |  | 95.00% CI of diff. |  | Adjusted P Value |
| CTRL vs. NASPM | 6.415 |  | -20.94 to 33.77 |  | 0.9248 |
| CTRL vs. Ketamine | -25.41 |  | -50.59 to -0.2344 |  | 0.0471 |
| CTRL vs. Ketamine + NASPM | 27.76 |  | 2.577 to 52.94 |  | 0.0253 |
| NASPM vs. Ketamine | -31.83 |  | -57.60 to -6.062 |  | 0.0097 |
| NASPM vs. Ketamine + NASPM | 21.34 |  | -4.426 to 47.11 |  | 0.1376 |
| Ketamine vs. Ketamine + NASPM | 53.17 |  | 29.73 to 76.61 |  | <0.0001 |

**Table 2.** Statistical analysis of the amplitudes of uEPSCs (V_hold_ = +60 mV) in hippocampal CA1 pyramidal neurons.

|  |  |  |  |  |  |
| --- | --- | --- | --- | --- | --- |
| Female (+60 mV) |  |  |  |  |  |
| ANOVA table | SS (Type III) | DF | MS | F (DFn, DFd) | P value |
| Interaction | 58023 | 1 | 58023 | F (1, 50) = 3.814 | P=0.0564 |
| Row Factor (Saline vs. Ket) | 78.38 | 1 | 78.38 | F (1, 50) = 0.005152 | P=0.9431 |
| Column Factor (Saline vs. NASPM) | 88106 | 1 | 88106 | F (1, 50) = 5.791 | P=0.0198 |
| Residual | 760699 | 50 | 15214 |  |  |
| Tukey's multiple comparisons test | Predicted (LS) mean diff. |  | 95.00% CI of diff. |  | Adjusted P Value |
| CTRL vs. NASPM | 15.24 |  | -111.0 to 141.5 |  | 0.9885 |
| CTRL vs. Ketamine | -68.02 |  | -196.6 to 60.56 |  | 0.5017 |
| CTRL vs. Ketamine + NASPM | 78.43 |  | -47.83 to 204.7 |  | 0.3602 |
| NASPM vs. Ketamine | -83.25 |  | -209.5 to 43.00 |  | 0.3082 |
| NASPM vs. Ketamine + NASPM | 63.19 |  | -60.70 to 187.1 |  | 0.5327 |
| Ketamine vs. Ketamine + NASPM | 146.4 |  | 20.19 to 272.7 |  | 0.0170 |
| Male (+60 mV) |  |  |  |  |  |
| ANOVA table | SS (Type III) | DF | MS | F (DFn, DFd) | P value |
| Interaction | 67842 | 1 | 67842 | F (1, 55) = 4.158 | P=0.0462 |
| Row Factor (Saline vs. Ket) | 4323 | 1 | 4323 | F (1, 55) = 0.2650 | P=0.6088 |
| Column Factor (Saline vs. NASPM) | 70148 | 1 | 70148 | F (1, 55) = 4.300 | P=0.0428 |
| Residual | 897324 | 55 | 16315 |  |  |
| Tukey's multiple comparisons test | Predicted (LS) mean diff. |  | 95.00% CI of diff. |  | Adjusted P Value |
| CTRL vs. NASPM | 1.157 |  | -134.3 to 136.6 |  | >0.9999 |
| CTRL vs. Ketamine | -85.98 |  | -210.7 to 38.70 |  | 0.2718 |
| CTRL vs. Ketamine + NASPM | 52.48 |  | -72.20 to 177.2 |  | 0.6818 |
| NASPM vs. Ketamine | -87.14 |  | -214.7 to 40.45 |  | 0.2799 |
| NASPM vs. Ketamine + NASPM | 51.32 |  | -76.27 to 178.9 |  | 0.7116 |
| Ketamine vs. Ketamine + NASPM | 138.5 |  | 22.39 to 254.5 |  | 0.0133 |

### CRS-induced social dysfunction is reversed by ketamine via CP-AMPARs

Repeated stress can induce the development of social dysfunction (daSilva et al., 2021). Importantly, it has been shown that an increase in hippocampal activity reverses stress-induced social dysfunction (Okuyama et al., 2016; Sun et al., 2020). We thus conducted a reciprocal social interaction test to examine whether chronic stress induced social dysfunction and if ketamine reversed this via CP-AMPARs. For chronic stress in mice, we used chronic restraint stress (CRS) because it has been widely used as a model of chronic psychoemotional stress to induce depression-and anxiety-like behaviors, learning and memory deficits, social dysfunction, and hippocampal neuronal damage in mice (Huang et al., 2015; Park, Seo, Lee, Shin, & Kang, 2018; Yun et al., 2010). After restraint stress, mice were returned to the home cage. Following 2-week CRS, we conducted the reciprocal social interaction test was carried out as shown in the experimental timeline in **Fig. 1**. First, we found that CRS significantly decreased the total number of reciprocal interactions (FEMALE: Naïve, 32.67 ± 8.41 and CRS, 16.83 ± 4.31, *p* = 0.0425. MALE: Naïve, 33.00 ± 10.94 and CRS, 16.77 ± 7.18, *p* = 0.0028) and total time of reciprocal interactions (FEMALE: Naïve, 16.28 ± 3.55 sec and CRS, 7.37 ± 4.83 sec, *p* = 0.0315. MALE: Naïve, 19.03 ± 6.56 sec and CRS, 9.02 ± 2.22 sec, *p* = 0.0026) in both female and male mice when compared to Naïve controls, an indication of social dysfunction (**Fig. 3 and Table 3-4**). This suggests that CRS significantly disrupted social behavior in animals. We then injected low-dose ketamine to CRS animals and examined whether ketamine reversed CRS-induced social dysfunction one hour after the treatment. We discovered that ketamine markedly increased the total number of reciprocal interactions (FEMALE: CRS + Ket, 44.29 ± 12.41, *p* < 0.0001. MALE: CRS + Ket, 33.25 ± 11.83, *p* = 0.0001) and total time of reciprocal interactions (FEMALE: CRS + Ket, 23.09 ± 6.29 sec, *p* < 0.0001. MALE: CRS + Ket, 17.71 ± 8.13 sec, *p* = 0.0009) in both female and male mice (**Fig. 3 and Table 3-4**). This suggests that subanesthetic low-dose ketamine can restore normal social behavior in chronically stressed animals. Finally, we tested whether these ketamine’s effects on social behavior were mediated by CP-AMPARs, we treated IEM-1460 (IEM), a BBB-permeable CP-AMPAR antagonist, with low-dose ketamine in CRS mice and carried out the social behavior test. We revealed that IEM treatment significantly reduced the total number of reciprocal interactions (FEMALE: CRS + Ket + IEM, 19.23 ± 9.08, *p* < 0.0001. MALE: CRS + Ket + IEM, 16.61 ± 7.33, *p* < 0.0001) and total time of reciprocal interactions (FEMALE: CRS + Ket + IEM, 11.60 ± 5.03 sec, *p* < 0.0001. MALE: CRS + Ket + IEM, 12.04 ± 4.38 sec, *p* = 0.0286) in both female and male mice when compared to ketamine-treated CRS mice (**Fig. 3 and Table 3-4**). These findings demonstrate that CRS impairs animals’ social behavior, which is reversed by low-dose ketamine treatment, and the ketamine’s effects are mediated by CP-AMPARs.

**Figure 3.**
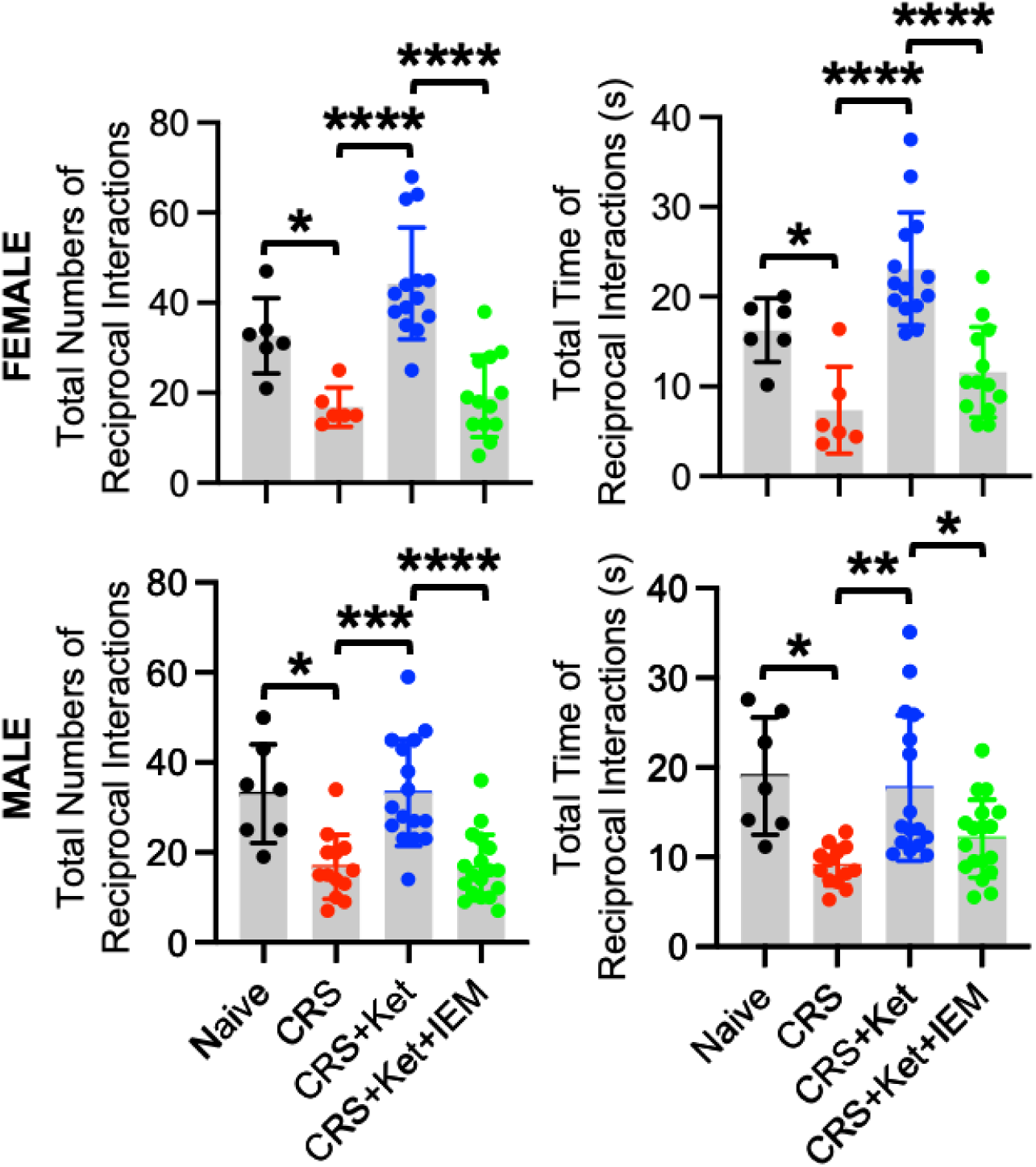
CRS-induced social dysfunction is reversed by ketamine via CP-AMPARs. Summary of total numbers and time of reciprocal interactions in each condition (n = number of animals. FEMALE: Naïve = 5, CRS = 6, CRS + Ket = 14, and CRS + Ket + IEM = 13. MALE: Naïve = 7, CRS = 12, CRS + Ket = 14, and CRS + Ket + IEM = 17). \**p* < 0.05, \*\**p* < 0.01, \*\*\**p* < 0.001, and \*\*\*\**p* < 0.0001, One-Way ANOVA, Tukey test. Ketamine: 5 mg/kg for females and 10 mg/kg for males. IEM-1460: 10 mg/kg for both females and males.

**Table 3.**
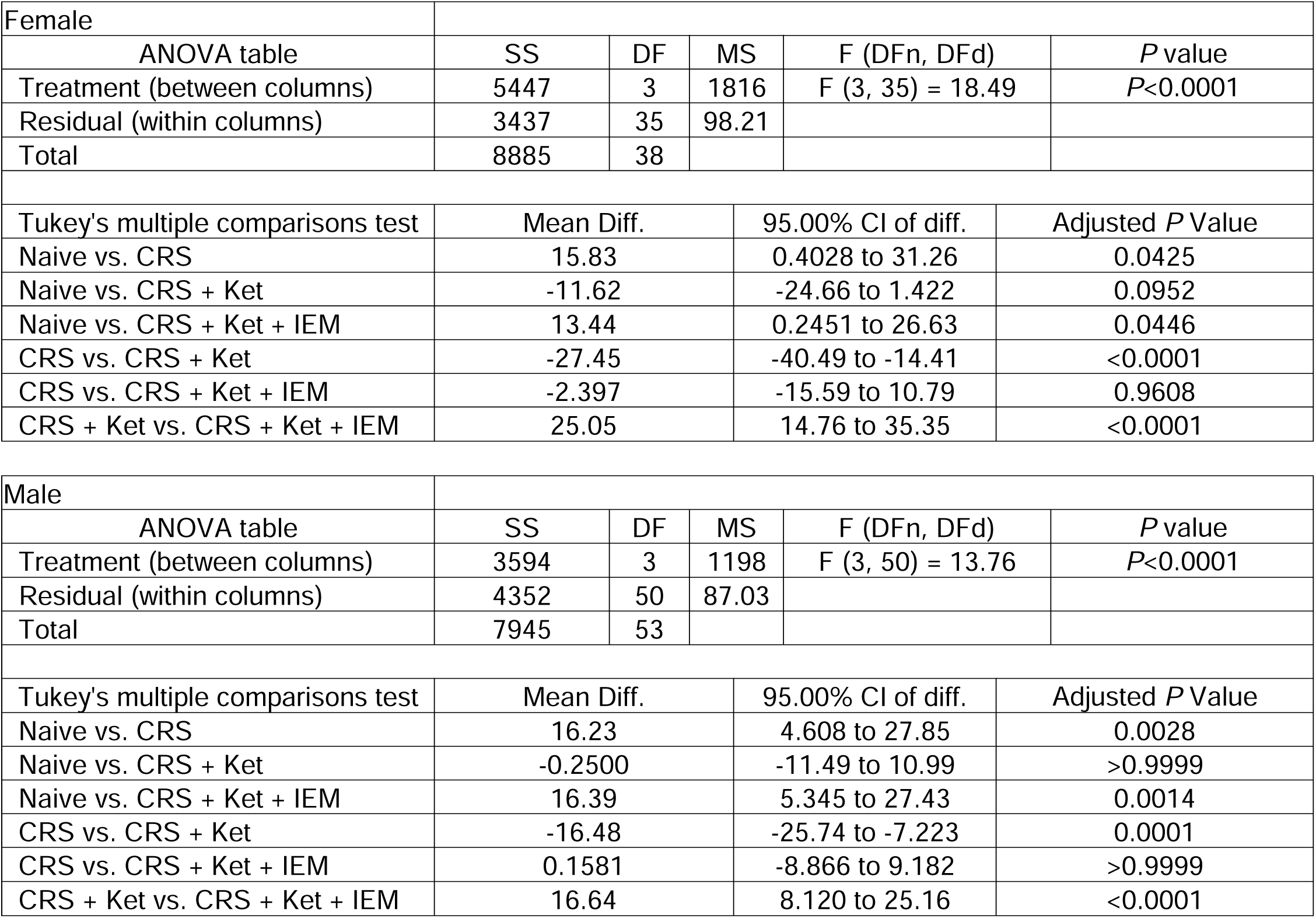
Statistical analysis of the total numbers of reciprocal interactions in the reciprocal social interaction test.

|  |  |  |  |  |  |
| --- | --- | --- | --- | --- | --- |
| Female |  |  |  |  |  |
| ANOVA table | SS | DF | MS | F (DFn, DFd) | P value |
| Treatment (between columns) | 5447 | 3 | 1816 | F (3, 35) = 18.49 | P<0.0001 |
| Residual (within columns) | 3437 | 35 | 98.21 |  |  |
| Total | 8885 | 38 |  |  |  |
| Tukey's multiple comparisons test | Mean Diff. |  | 95.00% CI of diff. |  | Adjusted P Value |
| Naive vs. CRS | 15.83 |  | 0.4028 to 31.26 |  | 0.0425 |
| Naive vs. CRS + Ket | -11.62 |  | -24.66 to 1.422 |  | 0.0952 |
| Naive vs. CRS + Ket + IEM | 13.44 |  | 0.2451 to 26.63 |  | 0.0446 |
| CRS vs. CRS + Ket | -27.45 |  | -40.49 to -14.41 |  | <0.0001 |
| CRS vs. CRS + Ket + IEM | -2.397 |  | -15.59 to 10.79 |  | 0.9608 |
| CRS + Ket vs. CRS + Ket + IEM | 25.05 |  | 14.76 to 35.35 |  | <0.0001 |
| Male |  |  |  |  |  |
| ANOVA table | SS | DF | MS | F (DFn, DFd) | P value |
| Treatment (between columns) | 3594 | 3 | 1198 | F (3, 50) = 13.76 | P<0.0001 |
| Residual (within columns) | 4352 | 50 | 87.03 |  |  |
| Total | 7945 | 53 |  |  |  |
| Tukey's multiple comparisons test | Mean Diff. |  | 95.00% CI of diff. |  | Adjusted P Value |
| Naive vs. CRS | 16.23 |  | 4.608 to 27.85 |  | 0.0028 |
| Naive vs. CRS + Ket | -0.2500 |  | -11.49 to 10.99 |  | >0.9999 |
| Naive vs. CRS + Ket + IEM | 16.39 |  | 5.345 to 27.43 |  | 0.0014 |
| CRS vs. CRS + Ket | -16.48 |  | -25.74 to -7.223 |  | 0.0001 |
| CRS vs. CRS + Ket + IEM | 0.1581 |  | -8.866 to 9.182 |  | >0.9999 |
| CRS + Ket vs. CRS + Ket + IEM | 16.64 |  | 8.120 to 25.16 |  | <0.0001 |

**Table 4.** Statistical analysis of the total time of reciprocal interactions in the reciprocal social interaction test.

|  |  |  |  |  |  |
| --- | --- | --- | --- | --- | --- |
| Female |  |  |  |  |  |
| ANOVA table | SS | DF | MS | F (DFn, DFd) | P value |
| Treatment (between columns) | 1401 | 3 | 466.9 | F (3, 35) = 16.40 | P<0.0001 |
| Residual (within columns) | 996.4 | 35 | 28.47 |  |  |
| Total | 2397 | 38 |  |  |  |
| Tukey's multiple comparisons test | Mean Diff. |  | 95.00% CI of diff. |  | Adjusted P Value |
| Naive vs. CRS | 8.917 |  | 0.6088 to 17.22 |  | 0.0315 |
| Naive vs. CRS + Ket | -6.802 |  | -13.82 to 0.2191 |  | 0.0605 |
| Naive vs. CRS + Ket + IEM | 4.683 |  | -2.419 to 11.79 |  | 0.3005 |
| CRS vs. CRS + Ket | -15.72 |  | -22.74 to -8.698 |  | <0.0001 |
| CRS vs. CRS + Ket + IEM | -4.233 |  | -11.34 to 2.869 |  | 0.3876 |
| CRS + Ket vs. CRS + Ket + IEM | 11.49 |  | 5.943 to 17.03 |  | <0.0001 |
| Male |  |  |  |  |  |
| ANOVA table | SS | DF | MS | F (DFn, DFd) | P value |
| Treatment (between columns) | 788.4 | 3 | 262.8 | F (3, 50) = 8.035 | P=0.0002 |
| Residual (within columns) | 1635 | 50 | 32.71 |  |  |
| Total | 2424 | 53 |  |  |  |
| Tukey's multiple comparisons test | Mean Diff. |  | 95.00% CI of diff. |  | Adjusted P Value |
| Naive vs. CRS | 10.01 |  | 2.888 to 17.14 |  | 0.0026 |
| Naive vs. CRS + Ket | 1.322 |  | -5.565 to 8.210 |  | 0.9563 |
| Naive vs. CRS + Ket + IEM | 6.990 |  | 0.2196 to 13.76 |  | 0.0406 |
| CRS vs. CRS + Ket | -8.691 |  | -14.37 to -3.016 |  | 0.0009 |
| CRS vs. CRS + Ket + IEM | -3.024 |  | -8.555 to 2.508 |  | 0.4734 |
| CRS + Ket vs. CRS + Ket + IEM | 5.667 |  | 0.4452 to 10.89 |  | 0.0286 |

### CRS-induced fear memory loss is reversed by ketamine via CP-AMPARs

Repeated stress can impair learning and memory (E. J. Kim & Kim, 2023). In fact, it has been shown that CRS disrupts hippocampus-dependent fear memory in mice (Yun et al., 2010). We thus examined whether low-dose ketamine reversed CRS-induced hippocampus-dependent fear memory loss via CP-AMPARs. To measure hippocampus-dependent fear memory, CFC was used (Tovote et al., 2015). As shown previously (Yun et al., 2010), we found that freezing was significantly lower in CRS female and male animals compared to naïve controls, an indication of fear memory loss (FEMALE: Naïve, 33.55 ± 11.50% and CRS, 15.24 ± 5.64%, *p* = 0.0026. MALE: Naïve, 26.45 ± 11.14% and CRS, 16.35 ± 7.64%, *p* = 0.0133) (**Fig. 4 and Table 5**). Next, low-dose ketamine was intraperitoneally injected to CRS female and male mice, and we carried out CFC to address the ketamine’s effects on fear memory. We discovered that ketamine treatment significantly increased freezing in CRS female and male mice (FEMALE: CRS + Ket, 28.16 ± 13.92%, *p* = 0.0454. MALE: CRS + Ket, 27.79 ± 11.50%, *p* = 0.0021) (**Fig. 4 and Table 5**). This suggests that subanesthetic low-dose ketamine can restore normal hippocampus-dependent fear memory in chronically stressed animals. To address that this ketamine’s effect was mediated by CP-AMPARs, we treated ketamine and IEM together in CRS mice. We revealed that blocking CP-AMPARs markedly reduced freezing in ketamine-treated CRS female and male animals compared to ketamine-treated CRS mice (FEMALE: CRS + Ket + IEM, 11.65 ± 4.45%, *p* = 0.0092.

**Figure 4.**
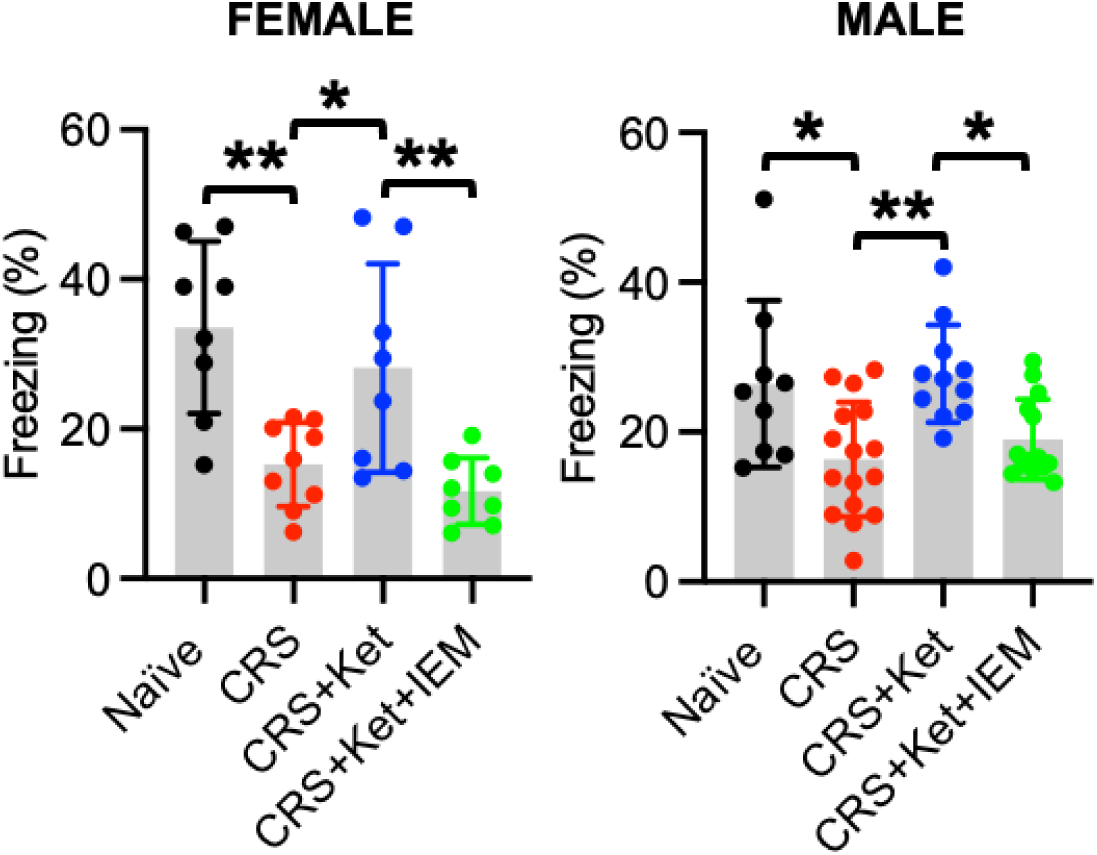
CRS-induced fear memory loss is reversed by ketamine via CP-AMPARs. Summary of fear memory in each condition (n = number of animals. FEMALE: Naïve = 8, CRS = 9, CRS + Ket = 8, and CRS + Ket + IEM = 8. MALE: Naïve = 9, CRS = 15, CRS + Ket = 11, and CRS + Ket + IEM = 14). \**p* < 0.05 and \*\**p* < 0.01, One-Way ANOVA, Tukey test. Ketamine: 5 mg/kg for females and 10 mg/kg for males. IEM-1460: 10 mg/kg for both females and males.

**Table 5.**
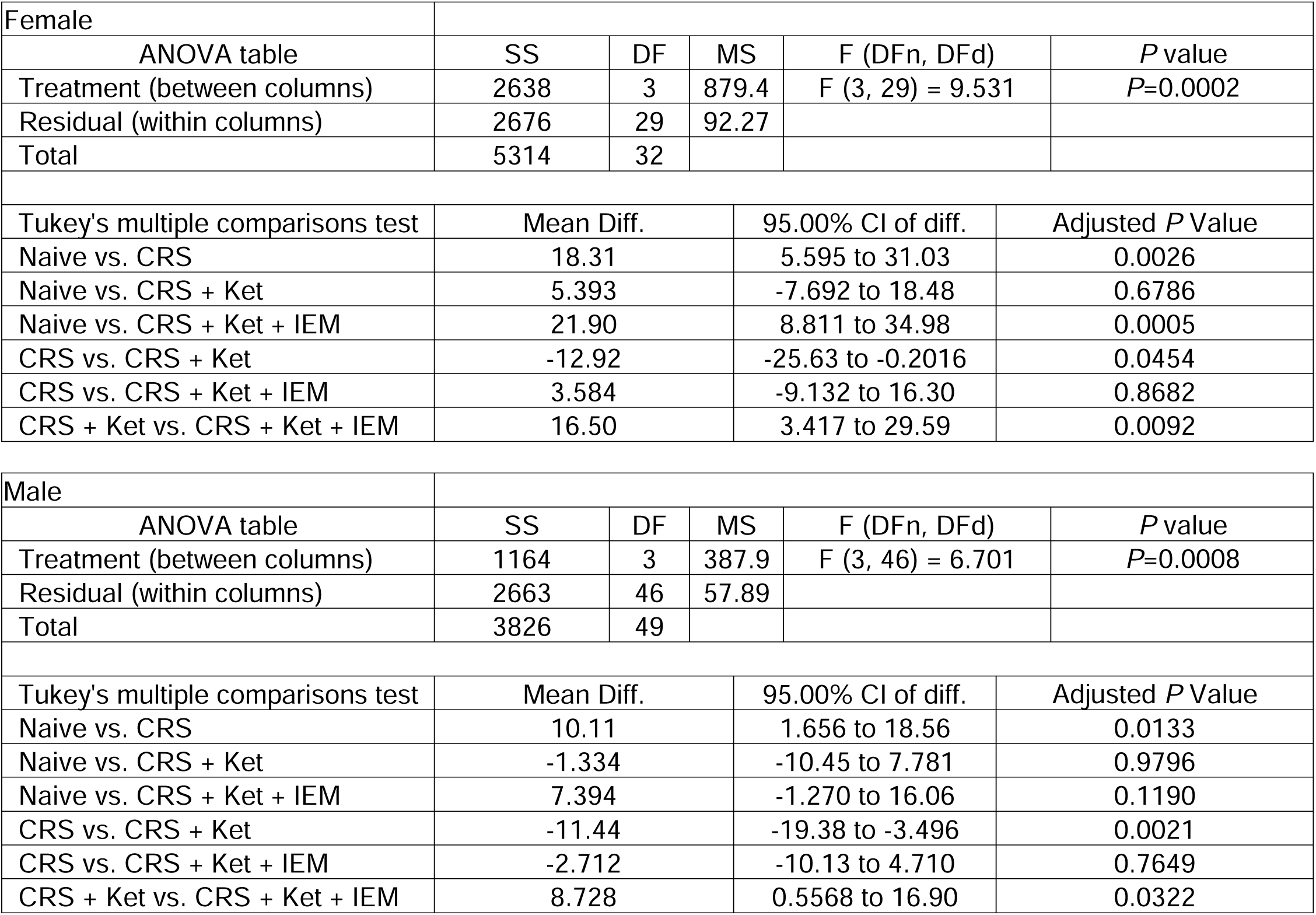
Statistical analysis of contextual fear conditioning.

|  |  |  |  |  |  |
| --- | --- | --- | --- | --- | --- |
| Female |  |  |  |  |  |
| ANOVA table | SS | DF | MS | F (DFn, DFd) | P value |
| Treatment (between columns) | 2638 | 3 | 879.4 | F (3, 29) = 9.531 | P=0.0002 |
| Residual (within columns) | 2676 | 29 | 92.27 |  |  |
| Total | 5314 | 32 |  |  |  |
| Tukey's multiple comparisons test | Mean Diff. |  | 95.00% CI of diff. |  | Adjusted P Value |
| Naive vs. CRS | 18.31 |  | 5.595 to 31.03 |  | 0.0026 |
| Naive vs. CRS + Ket | 5.393 |  | -7.692 to 18.48 |  | 0.6786 |
| Naive vs. CRS + Ket + IEM | 21.90 |  | 8.811 to 34.98 |  | 0.0005 |
| CRS vs. CRS + Ket | -12.92 |  | -25.63 to -0.2016 |  | 0.0454 |
| CRS vs. CRS + Ket + IEM | 3.584 |  | -9.132 to 16.30 |  | 0.8682 |
| CRS + Ket vs. CRS + Ket + IEM | 16.50 |  | 3.417 to 29.59 |  | 0.0092 |
| Male |  |  |  |  |  |
| ANOVA table | SS | DF | MS | F (DFn, DFd) | P value |
| Treatment (between columns) | 1164 | 3 | 387.9 | F (3, 46) = 6.701 | P=0.0008 |
| Residual (within columns) | 2663 | 46 | 57.89 |  |  |
| Total | 3826 | 49 |  |  |  |
| Tukey's multiple comparisons test | Mean Diff. |  | 95.00% CI of diff. |  | Adjusted P Value |
| Naive vs. CRS | 10.11 |  | 1.656 to 18.56 |  | 0.0133 |
| Naive vs. CRS + Ket | -1.334 |  | -10.45 to 7.781 |  | 0.9796 |
| Naive vs. CRS + Ket + IEM | 7.394 |  | -1.270 to 16.06 |  | 0.1190 |
| CRS vs. CRS + Ket | -11.44 |  | -19.38 to -3.496 |  | 0.0021 |
| CRS vs. CRS + Ket + IEM | -2.712 |  | -10.13 to 4.710 |  | 0.7649 |
| CRS + Ket vs. CRS + Ket + IEM | 8.728 |  | 0.5568 to 16.90 |  | 0.0322 |

MALE: CRS + Ket + IEM, 19.06 ± 5.34%, *p* = 0.0322) (**Fig. 4 and Table 5**). This data suggest that CRS disrupts hippocampus-dependent fear memory in animals, which is reversed by low-dose ketamine treatment, and the ketamine’s effect is dependent on CP-AMPARs.

### CRS-induced depression-like behavior is reversed by ketamine via CP-AMPARs

Repeated stress is a known trigger for depression in humans and depression-like behavior in animals (Richter-Levin & Xu, 2018; Tafet & Nemeroff, 2016). Indeed, CRS is known to induce depression-like behavior in rodents (Mao, Xu, & Yuan, 2022). Moreover, disruptions in AMPAR-mediated synaptic transmission in the hippocampus are associated with stress responses in animal models and individuals with depression (He et al., 2023). We thus employed TST to examine whether low-dose ketamine reversed CRS-induced depression-like behavior in mice via CP-AMPARs. As shown previously (Mao et al., 2022), CRS significantly elevated immobility in female and male mice when compared to Naïve controls, an indication of depression-like behavior (FEMALE: Naïve, 97.07 ± 42.05 sec and CRS, 168.80 ± 40.18 sec, *p* = 0.0130. MALE: Naïve, 78.80 ± 29.96 sec and CRS, 139.10 ± 36.04 sec, *p* = 0.0016) (**Fig. 5 and Table 6**). Importantly, low-dose ketamine treatment markedly decreased immobility in CRS female and male animals, an indication of antidepressant effects (FEMALE: CRS + Ket, 79.43 ± 35.91 sec, *p* = 0.0006. MALE: CRS + Ket, 90.25 ± 37.03 sec, *p* = 0.0066) (**Fig. 5 and Table 6**). We further discovered that immobility in IEM and ketamine-treated CRS mice was significantly higher than that ketamine-treated CRS mice (FEMALE: CRS + Ket + IEM, 131.50 ± 61.21 sec, *p* = 0.0472. MALE: CRS + Ket + IEM, 128.30 ± 31.56 sec, *p* = 0.0400) (**Fig. 5 and Table 6**). The data show that CRS induces depression-like behavior in animals, which is reversed by low-dose ketamine treatment via CP-AMPARs.

**Figure 5.**
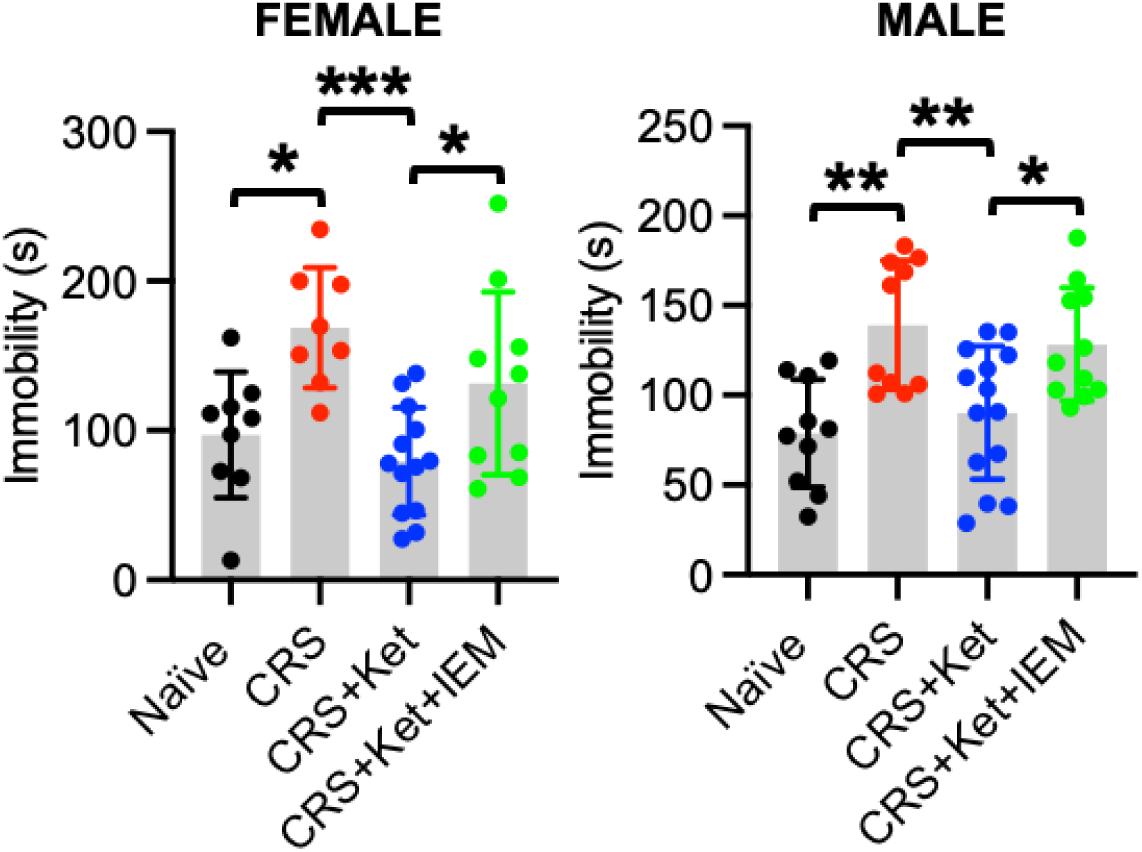
CRS-induced depression-like behavior is reversed by ketamine via CP-AMPARs. Summary of immobility in each condition (n = number of animals. FEMALE: Naïve = 9, CRS = 8, CRS + Ket = 13, and CRS + Ket + IEM = 10. MALE: Naïve = 10, CRS = 10, CRS + Ket = 14, and CRS + Ket + IEM = 11). \**p* < 0.05, \*\**p* < 0.01, and \*\*\**p* < 0.001, One-Way ANOVA, Tukey test. Ketamine: 5 mg/kg for females and 10 mg/kg for males. IEM-1460: 10 mg/kg for both females and males.

**Table 6.** Statistical analysis of the tail suspension test.

|  |  |  |  |  |  |
| --- | --- | --- | --- | --- | --- |
| Female |  |  |  |  |  |
| ANOVA table | SS | DF | MS | F (DFn, DFd) | P value |
| Treatment (between columns) | 45229 | 3 | 15076 | F (3, 36) = 7.272 | P=0.0006 |
| Residual (within columns) | 74637 | 36 | 2073 |  |  |
| Total | 119866 | 39 |  |  |  |
| Tukey's multiple comparisons test | Mean Diff. |  | 95.00% CI of diff. |  | Adjusted P Value |
| Naive vs. CRS | -71.77 |  | -131.4 to -12.18 |  | 0.0130 |
| Naive vs. CRS + Ket | 17.64 |  | -35.54 to 70.81 |  | 0.8084 |
| Naive vs. CRS + Ket + IEM | -34.42 |  | -90.77 to 21.92 |  | 0.3669 |
| CRS vs. CRS + Ket | 89.41 |  | 34.30 to 144.5 |  | 0.0006 |
| CRS vs. CRS + Ket + IEM | 37.35 |  | -20.82 to 95.52 |  | 0.3239 |
| CRS + Ket vs. CRS + Ket + IEM | -52.06 |  | -103.6 to -0.4780 |  | 0.0472 |
| Male |  |  |  |  |  |
| ANOVA table | SS | DF | MS | F (DFn, DFd) | P value |
| Treatment (between columns) | 27140 | 3 | 9047 | F (3, 41) = 7.801 | P=0.0003 |
| Residual (within columns) | 47549 | 41 | 1160 |  |  |
| Total | 74689 | 44 |  |  |  |
| Tukey's multiple comparisons test | Mean Diff. |  | 95.00% CI of diff. |  | Adjusted P Value |
| Naive vs. CRS | -60.31 |  | -101.1 to -19.53 |  | 0.0016 |
| Naive vs. CRS + Ket | -11.45 |  | -49.20 to 26.30 |  | 0.8484 |
| Naive vs. CRS + Ket + IEM | -49.48 |  | -89.32 to -9.640 |  | 0.0097 |
| CRS vs. CRS + Ket | 48.86 |  | 11.11 to 86.61 |  | 0.0066 |
| CRS vs. CRS + Ket + IEM | 10.83 |  | -29.01 to 50.67 |  | 0.8854 |
| CRS + Ket vs. CRS + Ket + IEM | -38.03 |  | -74.77 to -1.292 |  | 0.0400 |

### No abnormality in the open field test

As animals’ locomotor capacities can influence behavior, we carried out OFT to examine whether there were any abnormalities in animals’ locomotion in each condition. We found that there was no difference in total distance traveled, total time spent outside, and total time spent inside between the conditions (**Fig. 6 and Table 7**). Moreover, as time spent outside and inside can be interpreted as anxiety-like behavior (Zaytseva et al., 2023), normal behaviors in OFT indicate that animals used in our study had normal locomotor activity and no anxiety-like behavior. These findings suggest that behavioral alterations we described in the above are not due to changes in animals’ movement and anxiety levels.

**Figure 6.**
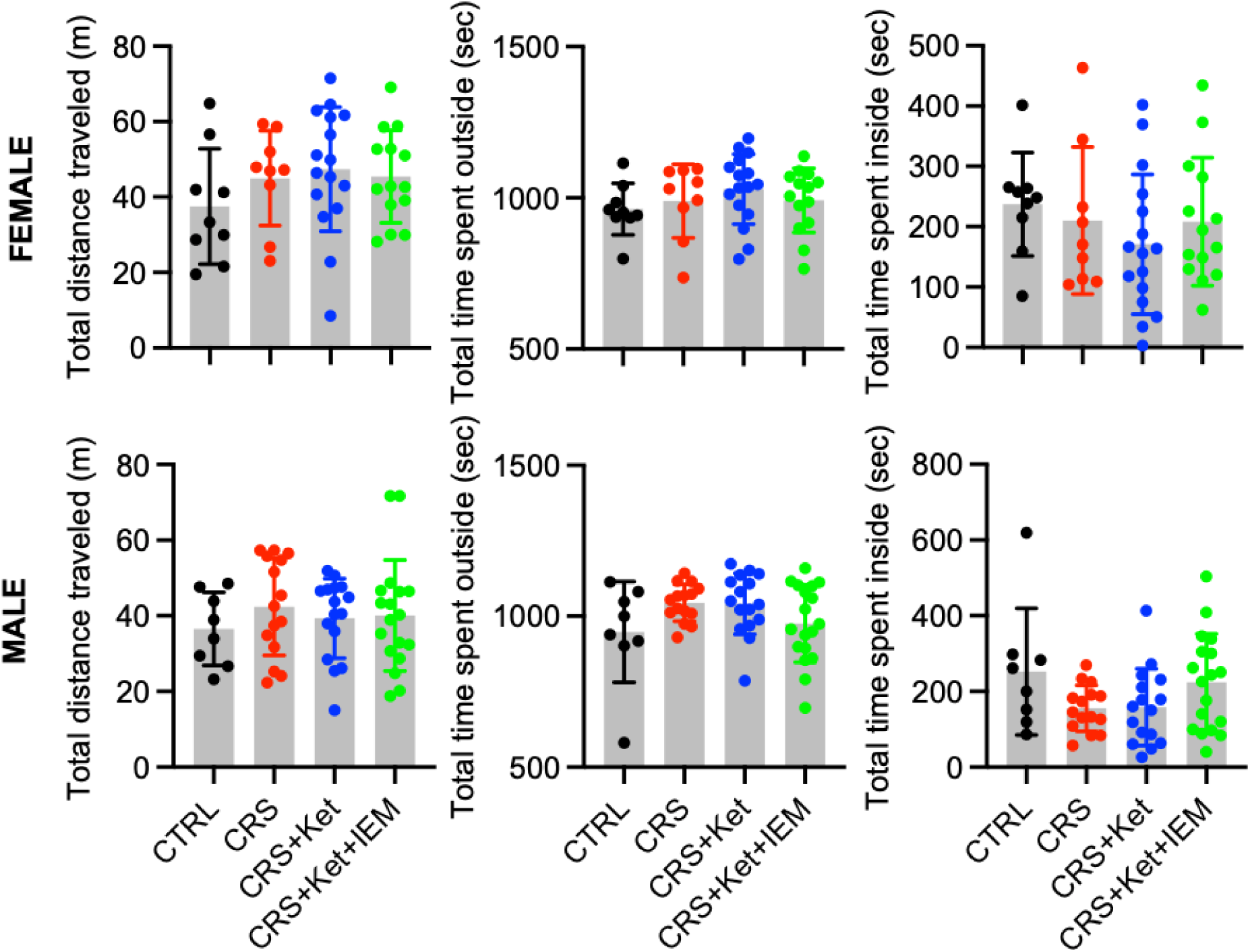
No abnormality in the open field test. Summary of total distance traveled, total time spent outside, and total time spent inside in each condition (n = number of animals. FEMALE: Naïve = 9, CRS = 9, CRS + Ket = 16, and CRS + Ket + IEM = 14. MALE: Naïve = 8, CRS = 15, CRS + Ket = 16, and CRS + Ket + IEM = 18). Ketamine: 5 mg/kg for females and 10 mg/kg for males. IEM-1460: 10 mg/kg for both females and males.

**Table 7.** Summary of the open field test.

|  | FEMALE |  |  |  | MALE |  |  |  |
| --- | --- | --- | --- | --- | --- | --- | --- | --- |
|  | CTRL | CRS | CRS +<br>Ket | CRS +<br>Ket + IEM | CTRL | CRS | CRS +<br>Ket | CRS +<br>Ket + IEM |
| Total distance traveled (m) | 37.52 ± 15.31 | 45.02 ± 12.62 | 47.37 ± 16.47 | 45.43 ± 12.24 | 36.55 ± 9.70 | 42.37 ± 12.86 | 39.34 ± 10.56 | 40.07 ± 14.65 |
| Total time spent outside (sec) | 962.8 ± 85.57 | 989.6 ± 122.0 | 1029 ± 115.9 | 991.8 ± 106.0 | 947.6 ± 167.2 | 1044 ± 60.74 | 1042 ± 101.2 | 975.9 ± 128.1 |
| Total time spent inside (sec) | 237.2 ± 85.57 | 210.4 ± 122.0 | 170.8 ± 115.9 | 208.2 ± 106.0 | 252.4 ± 167.2 | 155.6 ± 60.74 | 158.4 ± 101.2 | 224.1 ± 128.1 |

### Ketamine reverses the CRS-induced elevation of corticosterone levels, which is not dependent on CP-AMPARs

Chronic stress activates the hypothalamic-pituitary-adrenal axis, resulting in the release of glucocorticoid class stress hormones (McEwen, 2007; L. J. Phillips et al., 2006). Cortisol is the primary endogenous adrenal steroid in most mammals, including humans, whereas corticosterone is the primary adrenal corticosteroid in rodents (Raff, Sharma, & Nieman, 2014; Raff, Skelton, & Cowley, 1989; Usa et al., 2007; Yu et al., 2015). Chronic stress can keep these hormone levels high, which in turn alters brain functions (Burke, Sotiropoulos, & Waites, 2024; McEwen, 2007). In fact, CRS is known to increase corticosterone levels in animals, which is associated with stress-induced behavioral changes (Marin, Cruz, & Planeta, 2007). We thus examined whether low-dose ketamine reversed the CRS-induced elevation of corticosterone levels via CP-AMPARs. As shown previously (Tan, Wang, Chen, Long, & Zou, 2017), we found that CRS significantly increased serum corticosterone levels in both female and male animals when compared to naïve mice (FEMALE: Naïve, 831.1 ± 376.8 pg/ml and CRS, 2374 ± 1101 pg/ml, *p* = 0.0001. MALE: Naïve, 756.1 ± 210.6 pg/ml and CRS, 2140 ± 721.1 pg/ml, *p* = 0.0005) (**Fig. 7 and Table 8**). Ketamine is known to significantly decrease the CRS-induced increase in serum corticosterone levels in mice (Tan et al., 2017). Consistent with this finding, we discovered that low-dose ketamine significantly reduced serum corticosterone levels in CRS animals (FEMALE: CRS + Ket, 834.3 ± 372.0 pg/ml, *p* = 0.0001. MALE: CRS + Ket, 1089 ± 851.3 pg/ml, *p* = 0.0146) (**Fig. 7 and Table 8**). Finally, we tested whether the ketamine’s effects on corticosterone levels were mediated by CP-AMPARs. In contrast to behavioral assays, serum corticosterone levels of ketamine-treated CRS animals did not differ in the presence or absence of IEM (FEMALE: CRS + Ket + IEM, 1095 ± 481.9 pg/ml, *p* = 0.9247. MALE: CRS + Ket + IEM, 1082 ± 383.5 pg/ml, *p* > 0.9999) (**Fig. 7 and Table 8**). The data show that low-dose ketamine treatment reverses the CRS-induced increase in serum corticosterone levels, which is independent of CP-AMPARs.

**Figure 7.**
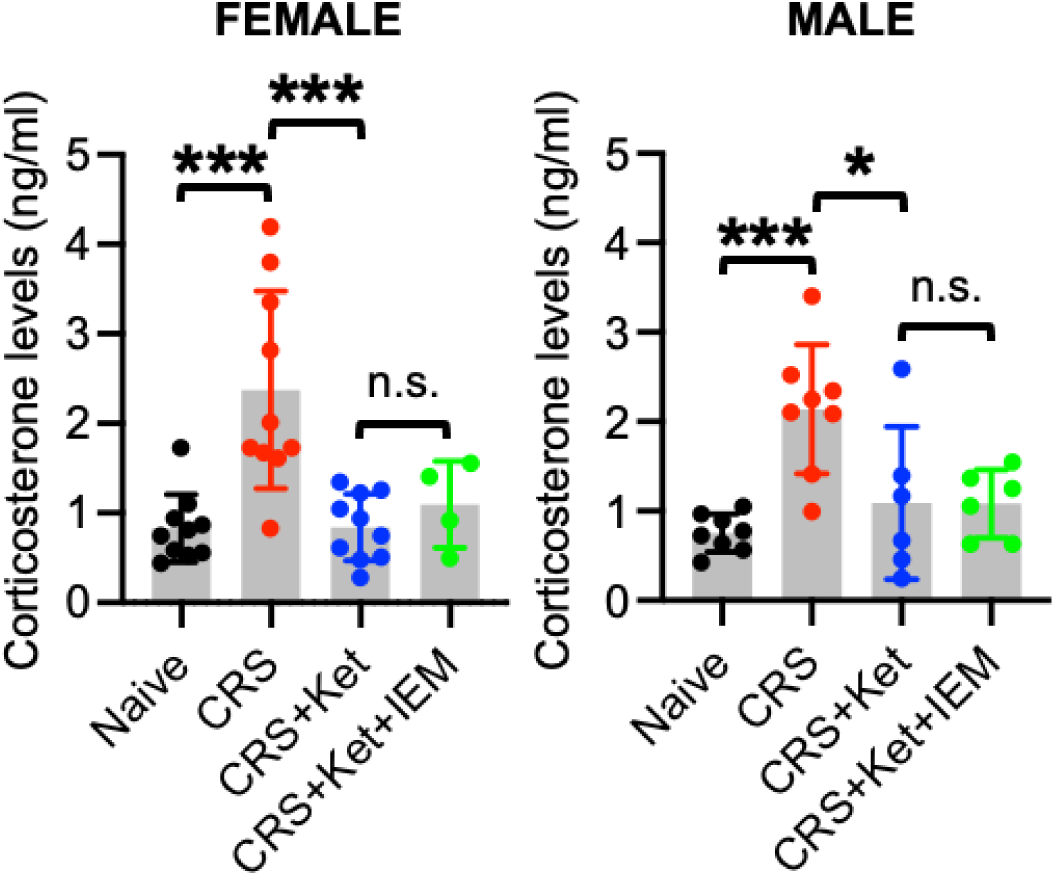
Ketamine reverses the CRS-induced elevation of corticosterone levels, which is not dependent on CP-AMPARs. Summary of serum corticosterone levels in each condition (n = number of animals with duplication. FEMALE: Naïve = 5, CRS = 5, CRS + Ket = 5, and CRS + Ket + IEM = 2. MALE: Naïve = 4, CRS = 4, CRS + Ket = 3, and CRS + Ket + IEM = 3). \**p* < 0.05 and \*\*\**p* < 0.001, One-Way ANOVA, Tukey test. n.s. – not significant. Ketamine: 5 mg/kg for females and 10 mg/kg for males. IEM-1460: 10 mg/kg for both females and males.

**Table 8.** Statistical analysis of corticosterone levels.

|  |  |  |  |  |  |
| --- | --- | --- | --- | --- | --- |
| Female |  |  |  |  |  |
| ANOVA table | SS | DF | MS | F (DFn, DFd) | *P* value |
| Treatment (between columns) | 15981461 | 3 | 5327154 | F (3, 30) = 11.31 | *P*<0.0001 |
| Residual (within columns) | 14132346 | 30 | 471078 |  |  |
| Total | 30113807 | 33 |  |  |  |
| Tukey's multiple comparisons test |  |  |  |  |  |
|  | Mean Diff. |  | 95.00% CI of diff. |  | Adjusted *P* Value |
| Naive vs. CRS | -1543 |  | -2378 to -708.6 |  | 0.0001 |
| Naive vs. CRS + Ket | -12.17 |  | -846.8 to 822.4 |  | >0.9999 |
| Naive vs. CRS + Ket + IEM | -264.2 |  | -1368 to 839.9 |  | 0.9145 |
| CRS vs. CRS + Ket | 1531 |  | 696.5 to 2366 |  | 0.0001 |
| CRS vs. CRS + Ket + IEM | 1279 |  | 175.0 to 2383 |  | 0.0183 |
| CRS + Ket vs. CRS + Ket + IEM | -252.0 |  | -1356 to 852.1 |  | 0.9247 |
| Male |  |  |  |  |  |
| ANOVA table | SS | DF | MS | F (DFn, DFd) | *P* value |
| Treatment (between columns) | 8562805 | 3 | 2854268 | F (3, 24) = 8.244 | *P*=0.0006 |
| Residual (within columns) | 8309607 | 24 | 346234 |  |  |
| Total | 16872412 | 27 |  |  |  |
| Tukey's multiple comparisons test |  |  |  |  |  |
|  | Mean Diff. |  | 95.00% CI of diff. |  | Adjusted *P* Value |
| Naive vs. CRS | -1384 |  | -2196 to -572.3 |  | 0.0005 |
| Naive vs. CRS + Ket | -332.8 |  | -1209 to 543.9 |  | 0.7239 |
| Naive vs. CRS + Ket + IEM | -325.8 |  | -1202 to 550.9 |  | 0.7366 |
| CRS vs. CRS + Ket | 1051 |  | 174.5 to 1928 |  | 0.0146 |
| CRS vs. CRS + Ket + IEM | 1058 |  | 181.5 to 1935 |  | 0.0139 |
| CRS + Ket vs. CRS + Ket + IEM | 7.000 |  | -930.2 to 944.2 |  | >0.9999 |

### Discussion and Conclusions Ketamine’s unknown mechanisms

It is challenging to understand how ketamine works due to its complex dose-dependent effects, with multiple pharmacologically active metabolites, and various mechanisms associated with the facilitation of synaptic plasticity (Kohtala, 2021). Importantly, the subanesthetic low-dose ketamine-induced increase of glutamatergic synaptic strength is presumed to underlie antidepressant effects in humans and animals (Aleksandrova et al., 2020; Kavalali & Monteggia, 2020; Miller et al., 2016). These actions of ketamine are likely mediated by enhancing AMPAR activity due to its nature of NMDAR antagonism (Aleksandrova et al., 2017). However, the entire role of AMPARs in the actions of ketamine remains undetermined. Additionally, treatment with ketamine is associated with psychomimetic side effects and risk of addiction, which prevent its broader application (Ng et al., 2010). Therefore, it is critical to identify the mechanisms of ketamine’s effects, particularly on glutamatergic synapses, to guide the development of safer fast-acting therapy.

### The role of CP-AMPARs in ketamine-induced antistress effects

Disruptions of AMPAR-mediated synaptic transmission in diverse cell types in the hippocampus are one of major biological changes in chronically stressed brains of animal models and individuals with depression (He et al., 2023). Importantly, recovery from GluA1-mediated synaptic dysfunction is thought to be the key target of ketamine’s antidepressant and antistress effects. Moreover, ketamine-induced recovery from disruptions in AMPAR-mediated synaptic transmission and behaviors in stressed animals is suggested to be associated with an increase in synaptic GluA1 expression in the hippocampus (Fischell et al., 2015; Li et al., 2011). In fact, we have discovered that low-dose ketamine rapidly induces the expression of CP-AMPARs in cultured hippocampal neurons, which is mediated by increasing GluA1 phosphorylation and its surface expression through a reduction of neuronal Ca^2+^ and Ca^2+^-dependent phosphatase calcineurin (CaN) activity (Zaytseva et al., 2023) (**Fig. 8**). In particular, GluA1-containing CP-AMPARs have higher single channel conductance, which contributes to synaptic plasticity (Aoto et al., 2008; Goel & Lee, 2007; Goel et al., 2011; S. Kim et al., 2015; S. Kim & Ziff, 2014; H. K. Lee, 2012; Purkey & Dell’Acqua, 2020; Sanderson et al., 2016; Sanderson et al., 2018; Thiagarajan et al., 2005). We have indeed shown that ketamine-induced CP-AMPAR expression enhances glutamatergic synaptic strength and chemically-induced long-term potentiation (cLTP) in cultured neurons (Zaytseva et al., 2023). cLTP is shown to rely on a rise in postsynaptic Ca^2+^ concentration, specifically through the activation of NMDARs (Molnar, 2011). Therefore, when ketamine is treated, CP-AMPAR-mediated Ca^2+^ influx may replace NMDAR-dependent Ca^2+^ signaling. We further show that subanesthetic low-dose ketamine decreases anxiety-and depression-like behaviors in naïve animals (Zaytseva et al., 2023). Importantly, an increase in hippocampal activity is known to improve stress-induced negative behaviors in mice (Aleksandrova et al., 2020; Bird & Burgess, 2008; Grieco et al., 2022; Kavalali & Monteggia, 2020; Miller et al., 2016; Okuyama et al., 2016; Sun et al., 2020; Tovote et al., 2015), which further supports our findings. However, the mechanisms of ketamine’s protective effects against stress-induced disorders are not completely understood. Here, our electrophysiological data demonstrate that low-dose ketamine induces synaptic insertion of CP-AMPARs in pyramidal neurons of the hippocampus CA1 area.

**Figure 8.**
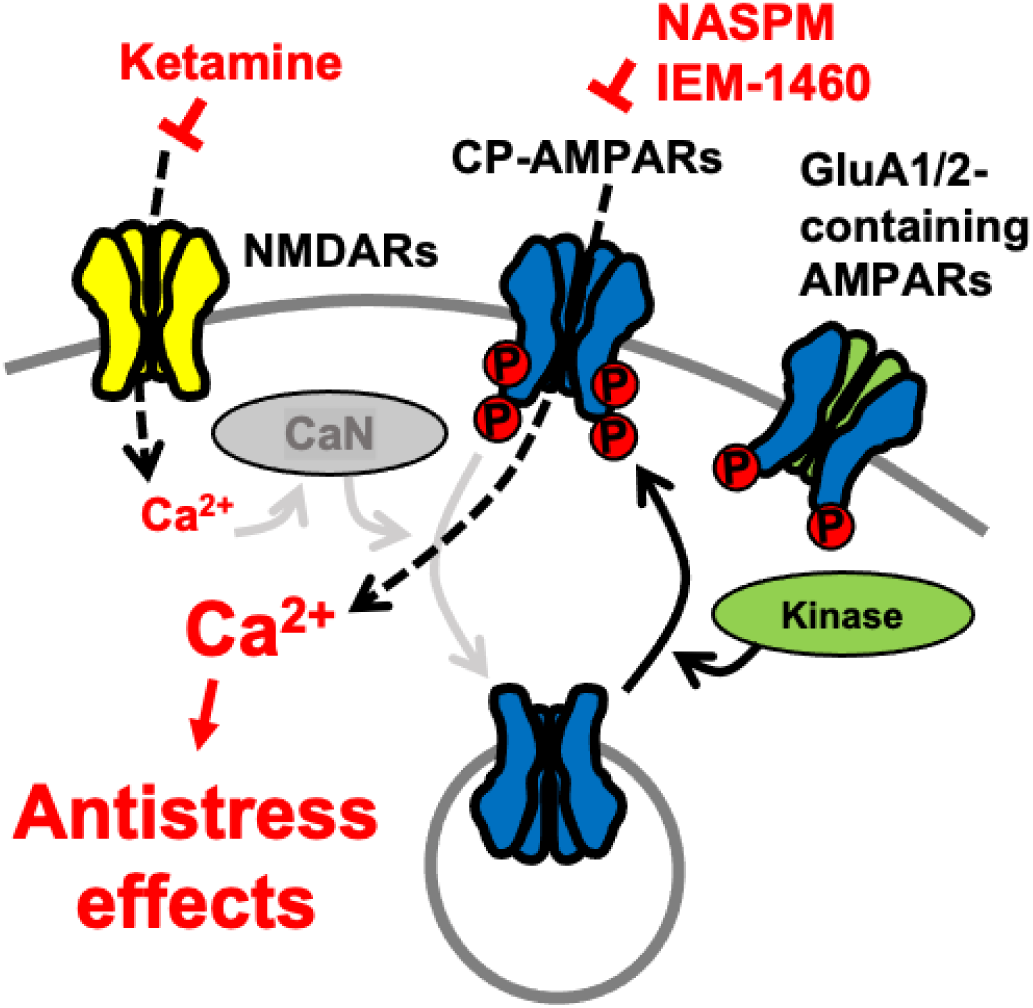
A schematic model of ketamine’s antistress effects. Because ketamine is a noncompetitive NMDAR antagonist, a therapeutic dose is enough to block NMDAR-mediated Ca^2+^ influx in excitatory synapses. This can lower the activity of calcineurin (CaN), a Ca^2+^-dependent phosphatase that dephosphorylates the AMPA receptor (AMPAR) subunit GluA1, and promote GluA1 phosphorylation, resulting in increased synaptic insertion of GluA2-lacking, GluA1-containing Ca^2+^-permeable AMPARs (CP-AMPARs). When ketamine is administered, CP-AMPAR-mediated Ca^2+^ influx may replace NMDAR-dependent Ca^2+^ signaling. This increases neural plasticity, which leads to antistress effects in animals. Therefore, inhibition of CP-AMPARs using antagonists such as NASPM and IEM-1460 prevents the ketamine’s effects.

Additionally, the current study reveals that subanesthetic low-dose ketamine can rapidly reverse social dysfunction, fear memory loss, and depression-like behavior caused by CRS in mice via CP-AMPARs (**Fig.8**). This suggests that low-dose ketamine induces synaptic insertion of CP-AMPARs in the hippocampus, which in turn enhances synaptic strength to reverse hippocampus-dependent behavioral dysfunctions in chronically stressed animals.

### The ketamine’s rapid onset and long-lasting effects

Given its brief half-life (3 hours in humans and 13 min in mice), the ketamine’s rapid onset and long-lasting effects (which persist for days in humans and 24 hours in mice) are not fully understood by synaptic mechanisms (Silva & Proulx, 2024). One potential mechanism underlying the ketamine’s rapid onset and long-lasting effects is that ketamine induces neural plasticity (Wu et al., 2021). However, still unclear, is when and how the plasticity is induced. Specifically, molecular and cellular processes of how ketamine induces neural plasticity remain to be elucidated. In addition to strengthening excitatory synaptic drive, low-dose ketamine can enhance neuronal Ca^2+^ signaling in the hippocampus (Zucker, 1999), possibly contributing to the ketamine’s rapid onset and long-lasting effects. Although the relationship between ketamine-triggered alteration in Ca^2+^ signaling and functional consequences for downstream processes regulating key aspects of neuronal function is known to be important for the antidepressant and antistress effects (Lisek, Zylinska, & Boczek, 2020), how ketamine enhances synaptic Ca^2+^ signaling is not fully understood. Given that CP-AMPARs are known to contribute to various forms of synaptic plasticity (Aoto et al., 2008; Cull-Candy & Farrant, 2021; Goel & Lee, 2007; Goel et al., 2011; Guire, Oh, Soderling, & Derkach, 2008; S. Kim et al., 2015; S. Kim & Ziff, 2014; H. K. Lee, 2012; Man, 2011; Purkey & Dell’Acqua, 2020; Sanderson et al., 2016; Sanderson et al., 2018; Thiagarajan et al., 2005), CP-AMPAR-mediated neural plasticity can facilitate the ketamine’s effects. Therefore, our findings in which low-dose ketamine rapidly induces hippocampal CP-AMPARs can be significant because ketamine can enhance not only synaptic transmission but also synaptic Ca^2+^ signaling in response to partial antagonism of NMDARs (**Fig. 8**). Therefore, the ketamine-induced synaptic insertion of CP-AMPARs can increase glutamatergic activity and compensate for reduced NMDAR inhibition-mediated synaptic Ca^2+^ signaling enabling the ketamine’s rapid onset and long-lasting effects.

### Sex differences in ketamine’s effects

Our work shows an important finding: female mice have higher ketamine antidepressant responses than male mice, as seen by synaptic insertion of CP-AMPARs following a lower ketamine dosage, which induces the antistress effects. Several studies including both male and female animals consistently demonstrate that females are more sensitive to ketamine (Carrier & Kabbaj, 2013; Dossat, Wright, Strong, & Kabbaj, 2018; Franceschelli, Sens, Herchick, Thelen, & Pitychoutis, 2015; Zanos et al., 2016). These studies collectively suggest that hormonal differences and metabolic factors contribute to the heightened sensitivity of females to ketamine’s effects at lower doses. In addition, the distinct pharmacokinetics of ketamine in the animals’ brain and plasma could be one reason for the increased ketamine’s antistress reactions in female mice (Saland & Kabbaj, 2018). Over the course of three hours after treatment, female rats’ medial prefrontal cortex and hippocampus contain higher levels of ketamine and norketamine, a metabolite of ketamine, than do male rats (Saland & Kabbaj, 2018). The study also shows that female rats’ slower clearance rates and longer half-lives result in higher post-treatment effects of ketamine and its metabolites (Saland & Kabbaj, 2018).

Unfortunately, the potential sex differences in response to ketamine have been particularly understudied at this time, which is an important area for the future investigation in order to develop tailored treatment and dosing in clinics.

### Multiple mechanisms underlying ketamine’s effects

The mechanisms underlying ketamine’s antidepressant effects are not fully understood, although multiple ideas have been proposed. These hypotheses include *1)* direct NMDAR inhibition-mediated mechanism in excitatory neurons (Chen et al., 2024; Kavalali & Monteggia, 2020; S. Ma et al., 2023; Miller et al., 2016; Y. Yang et al., 2018; Zaytseva et al., 2023); *2)* NMDAR inhibition-independent, ketamine metabolite-dependent mechanism (Carrier & Kabbaj, 2013; Franceschelli et al., 2015; Zanos et al., 2016); and *3)* NMDAR inhibition-induced disinhibition-and BDNF-induced protein synthesis-dependent mechanism (Deyama & Duman, 2020; Gerhard et al., 2020). Low-dose MK-801, a NMDAR antagonist, has shown to improve depression-like behavior and working memory in mice (Wesierska, Duda, & Dockery, 2013; B. Yang, Ren, Ma, Chen, & Hashimoto, 2016). Conversely, others suggest that the metabolism of ketamine to (2R,6R)-hydroxynorketamine (HNK), devoid of NMDAR inhibition properties, is important for the antidepressant effect (Carrier & Kabbaj, 2013; Franceschelli et al., 2015; Zanos et al., 2016). Although multiple studies including our own support the NMDAR inhibition-mediated mechanisms, it is important to investigate how our current findings integrate or act independently from other described mechanisms.

### Brain region-specific actions of ketamine

Research identifying the mechanisms of the antidepressant effects of ketamine has largely focused on the hippocampus, since it is known to be dysfunctional in depressive disorders (Carreno et al., 2016; Duman, Aghajanian, Sanacora, & Krystal, 2016; Moda-Sava et al., 2019; Zanos et al., 2016). More recently, other areas of the brain have been investigated, including the lateral habenula (Chen et al., 2024; S. Ma et al., 2023; Y. Yang et al., 2018). A recent study shows that in CRS animals, ketamine selectively inhibited NMDARs in lateral habenula neurons, but not in hippocampal pyramidal neurons. This suggests that the lateral habenula-NMDAR inhibition–dependent mechanism likely occurs more upstream in the cascade of ketamine signaling *in vivo* (Chen et al., 2024). In contrast, another recent study using a systematic and unbiased mapping approach that provides a comprehensive coverage of all brain regions discovers that ketamine preferably targets the hippocampus (Pasha A. Davoudian, Shao, & Kwan, 2022). The hippocampus is a crucial brain area that governs learning, memory, social interaction, and depression (Bird & Burgess, 2008; Campbell & Macqueen, 2004; Fortin, Agster, & Eichenbaum, 2002; Ghasemi, Navidhamidi, Rezaei, Azizikia, & Mehranfard, 2022; Jarrard, 1993; Xu et al., 2021). Research also demonstrates that an increase in hippocampal activity can be protective against stress (Aleksandrova et al., 2020; Bird & Burgess, 2008; Grieco et al., 2022; Kavalali & Monteggia, 2020; Miller et al., 2016; Okuyama et al., 2016; Sun et al., 2020; Tovote et al., 2015). Importantly, one of the most vulnerable targets of stress is hippocampus, because it abundantly expresses both glucocorticoid and mineralocorticoid receptors (Smith, 1996; Uno, Tarara, Else, Suleman, & Sapolsky, 1989). Moreover, stress changes hippocampal neural activity and synaptic plasticity, activating hippocampal glucocorticoid receptors, and decreasing neuronal cell survival and neurogenesis (Egeland, Zunszain, & Pariante, 2015; J. J. Kim & Diamond, 2002; Ryu et al., 2016). This suggests that the hippocampus is the important brain region responsible for ketamine’s effects. Since ketamine is delivered systemically in most studies, many regions in the brain can potentially be responsive to ketamine. Therefore, it is crucial to identify the crosstalk from the hippocampus to other brain regions including the lateral habenula for better understanding of the ketamine’s therapeutic effects.

### Potential role of ketamine’s anti-inflammatory properties

Chronic stress is known to increase the release of corticosterone in animals (McEwen, 2007; L. J. Phillips et al., 2006). In fact, we discover that ketamine significantly reduces serum corticosterone levels in CRS mice, which is consistent with the previous finding (Tan et al., 2017). In contrast to ketamine-induced protective effects against CRS-induced behavioral changes, this effect is not dependent on CP-AMPAR expression. Interestingly, chronic stress can induce disorders through increased pre-inflammatory cytokines, reactive microglia numbers, and up-regulated regulatory molecules (Liu, Wang, & Jiang, 2017). Moreover, chronic inflammatory responses directly affect brain structure and function, resulting in pathological behaviors in animals (McEwen, Nasca, & Gray, 2016). Importantly, ketamine is known to be associated with anti-inflammatory effects in humans (Dale, Somogyi, Li, Sullivan, & Shavit, 2012; Nikkheslat, 2021) and animals (Ommati et al., 2023; Spencer et al., 2022; N. Wang et al., 2015), suggesting that its ability to reduce the inflammation may play a role in its effects on ketamine’s protective effects against stress. Therefore, it is possible that the mechanism underlying the ketamine-induced decrease in stress hormones following chronic stress can be mediated by the hypothalamic-pituitary-adrenal axis rather than the hippocampus. It is thus possible that hypothalamic neurons may not express CP-AMPARs in response to ketamine treatment. In fact, there are multiple mechanisms proposed to explain the ketamine’s anti-inflammatory effects, including the involvement of the gut microbiome (Getachew et al., 2018; Zhang et al., 2017) and priming of Type 1 T helper (Th1)-type immune response (Ohta, Ohashi, & Fujino, 2009). However, clinical relevance of ketamine-induced inflammatory modulation is still unclear. Therefore, further studies are needed to identify the precise mechanisms of its anti-inflammatory properties.

## Conclusions

The current study addresses how chronic stress and ketamine affect hippocampal synaptic activity and hippocampus-dependent behaviors, providing a new evidence-based idea that NMDAR inhibition by ketamine leads to the antistress effects through synaptic insertion of CP-AMPARs. Therefore, our work advances current understandings on the mechanisms of ketamine’s antistress effects to guide the developments of new therapies.

## Acknowledgments

We thank members of the Kim laboratory for their generous support. This work is supported by Student Experiential Learning Grants and College Research Council Shared Research Program from College of Veterinary Medicine and Biomedical Sciences, Colorado State University and a research grant from Korea Ginseng Corporation.

